# Brain-wide mapping of oligodendrocyte organization and oligodendrogenesis across the murine lifespan

**DOI:** 10.1101/2024.09.06.611254

**Authors:** Yu Kang T. Xu, Abigail Bush, Ephraim Musheyev, Anya A. Kim, Sen Zhang, Jaime Eugenin von Bernhardi, Jeremias Sulam, Dwight E. Bergles

## Abstract

Insulating sheaths of myelin accelerate neuronal signaling in complex networks of the mammalian brain. In the CNS, myelin sheaths are exclusively produced by oligodendrocytes, which continue to be generated throughout life to change patterns of myelination. However, a brain-wide analysis of oligodendrocyte dynamics across the lifespan has not been performed. We developed a rapid, robust cellular mapping pipeline involving tissue clearing, lightsheet microscopy, atlas alignment, and automated segmentation to define the location of all oligodendrocytes in the mouse brain. This analysis demonstrated the remarkable consistency of oligodendrocyte patterns between hemispheres, individuals, and sexes, and established that oligodendrocyte maps estimate myelin coverage. We trained a vision transformer to identify newly generated oligodendrocytes from millions of mature cells, highlighting age- and region-specific differences in oligodendrogenesis, and revealing areas of enhanced oligodendrocyte resilience and regenerative capacity following demyelination, demonstrating the utility of this pipeline for uncovering brain-wide oligodendrocyte dynamics in health and disease.

## INTRODUCTION

Many axons in the mammalian CNS are ensheathed by myelin, a multilamellar membrane that acts as insulation to accelerate the propagation of action potentials^1^. The evolutionary development of myelin was crucial for expanding the size and complexity of the vertebrate nervous system^2^, allowing information to be transmitted rapidly in expansive, highly interconnected neural networks. In the brain, myelin is exclusively produced by oligodendrocytes, each of which extends cytoplasmic processes into the surrounding neuropil to form up to 60 discrete myelin segments (internodes) around axons in a local area^3,4^. The generation of oligodendrocytes (oligodendrogenesis) occurs exclusively through the differentiation of oligodendrocyte precursor cells (OPCs), which in mouse and human begins during late stages of development and continues into adulthood^5–9^, resulting in a progressive increase in myelin content and varying patterns of myelin as we age^10–12^. Both oligodendrogenesis and myelination are influenced by life experiences, such as social interaction^13,14^, sensory input^15^, and learning^16–19^, and vary significantly between neuron types, due to differences in axon diameter, levels of activity, and molecular diversity^20–23^. As a result, the distribution of oligodendrocytes and myelin is not uniform, but varies markedly between brain regions, most notably delineating white from gray matter, but also exhibiting extensive heterogeneity between discrete cortical and subcortical areas^20^. Mapping these patterns of myelination across the brain would help reveal how myelin is used to endow brain circuits with distinct functional characteristics. Moreover, comparing regional gradients in myelin patterning would help to define the molecular mechanisms responsible for these diverse patterns, and determine how loss of oligodendrocytes, as a result of injury, aging or disease, can impact sensory, motor and cognitive processes.

Brain-wide maps of myelin distribution have been generated through magnetic resonance imaging methods, which detect the restricted diffusion of water within compact myelin^24,25^, as well as susceptibility tensor imaging, which visualizes the anisotropic magnetic susceptibility of myelin sheaths^26^. Although these methods allow images to be taken of the living brain and can be used to follow progressive changes in myelin, they are very low resolution and less sensitive in gray matter regions where the density of myelin is lower and their orientation less uniform. Moreover, as they do not visualize myelin directly, they depend on assumptions about the origin of signals that may not hold in all circumstances^27^. Diverse methods exist to obtain high resolution images of myelin within brain tissue, ranging from histochemical methods that react with lipids (Black Gold, Sudan Black), immunolabeling against myelin proteins (MBP, MOG), or label free imaging that exploit the highly ordered, dense packing nature of myelin, including third harmonic generation^28^, coherent anti-Stokes Raman scattering (CARS)^29^, and spectral confocal reflectance microscopy (SCoRe)^30^. However, such approaches are difficult to apply at scale to obtain quantitative information about regional myelin patterns or compare across different samples, due to labeling inhomogeneity or inherent non-linearities in signal generation. At the highest resolution, serial electron microscopy (EM) provides unambiguous demarcation of myelin, that when combined with automated segmentation algorithms, can provide complete reconstructions of oligodendrocytes with their complement of myelin sheaths^31^. However, serial EM requires expensive specialized instrumentation, provides information from only small regions, and cannot presently be applied at scale to explore dynamic changes requiring analysis of samples from multiple individuals.

Recent advances combining transgenic expression of fluorescent proteins with whole tissue laser microscopy have made it possible to create high resolution maps of cell structure and vascular organization within the intact mouse brain^32–34^. By combining delipidation methods (tissue clearing) to reduce photon scattering, with single plane illumination (lightsheet) microscopy for rapid imaging of large volumes, it is possible to obtain high resolution images of fluorescent structures throughout whole tissues such as the brain. Moreover, advanced artificial intelligence (AI)-based analysis methods, such as deep learning, can be used to develop computational models to automatically identify features of interest, extracting quantitative information from millions of cells in 3D volumes that would otherwise not be quantifiable with manual human effort^35,36^. However, methodologies tailored for visualizing oligodendrocytes brain-wide have not yet been developed and no analytical models currently exist to consistently identify and segment these cells across different samples.

Here, we defined the regional distribution of oligodendrocytes in the mouse brain across the lifespan by imaging endogenous fluorescence in transgenic mice that express EGFP exclusively in oligodendrocytes (*Mobp-EGFP*, GENSAT^37,38^), overcoming the need to directly detect myelin. We adapted aqueous clearing methods that rendered brain tissue optically transparent, while preserving endogenous fluorescence, and collected serial 3D fluorescent images of the brain using lightsheet microscopy. Stitching and registration algorithms were optimized to align terabyte-sized volumes to a common coordinate framework for spatial analysis, and trained convolutional neural network models were developed to reliably determine the position, size and shape of millions of oligodendrocytes within the murine brain. We further employed vision transformers^39^ to identify rare cells, oligodendrocytes that had been recently generated, providing a snapshot of oligodendrogenesis without the need for transgenic or histological demarcation.

Using this clearing-imaging-analysis pipeline, we defined the regional changes in oligodendrocyte density that occur across most of the mouse lifespan, from two months to more than two years. Due to the speed and reliability of this pipeline, it was possible to extract quantitative information from 30 brains across several experimental cohorts, define inter-individual variance in oligodendrocyte patterning, and explore the effects of age, sex, strain, and injury on oligodendrocyte distribution. This analysis revealed that the remarkable heterogeneity in oligodendrocyte distribution between brain regions, which varied over four orders of magnitude, was highly consistent between hemispheres, individuals, and sex. Tracking these density changes across the murine lifespan uncovered brain-wide temporal shifts in oligodendrocyte abundance, which increased proportionally between discrete brain regions to preserve their relative differences. We developed additional analysis methods to relate myelin territory to oligodendrocyte abundance, showing that the oligodendrocyte soma analysis can be used as a proxy for assessing myelination. Finally, we used a demyelination model (cuprizone)^4^ to determine brain-wide differences in oligodendrocyte vulnerability and regeneration, revealing regional heterogeneities that may impact myelin repair in diseases like multiple sclerosis (MS). These segmented datasets of oligodendrocytes, oligodendrogenesis, and myelin territory, provide new insight into the dynamic patterning of oligodendrocytes in the mammalian brain, and our new analysis tools provide the means to study how this process is influenced by life experience, aging, and disease.

## RESULTS

### Aqueous clearing enables visualization of oligodendrocytes brain-wide

Oligodendrocytes are present in almost all regions of the brain and generate remarkably diverse myelination patterns. However, brain-wide comparative analysis between regions has been difficult, as the high density of myelin protein epitopes, particularly in white matter, reduces the penetration of antibodies in tissue and results in labeling inhomogeneities when staining whole brains (**Figure S1A**). To enable whole-brain visualization of oligodendrocytes using lightsheet microscopy, we developed a tissue clearing pipeline that preserves endogenous fluorescence (see Methods). This method allowed quantitative assessments of oligodendrocyte and myelin sheath distribution in *Mobp-EGFP* mice^37^ (**Figure 1A**), in which all ASPA immunoreactive (+) oligodendrocytes express EGFP, a pattern that was preserved in gray matter (GM) and white matter (WM), between mouse strains, and across the lifespan (percentage of EGFP+ cells that are also ASPA+: >99% for all groups, n=4 volumes per condition, **Figure 1B,C**). We found that many established clearing techniques (e.g. iDISCO^40^) could not preserve endogenous fluorescence, often quenching it entirely (**Figure S1B**), or were unable to achieve sufficient clarity to image deeper brain regions (**Figure S1C**). To establish a reproducible and accessible clearing approach, we combined aspects of three clearing methods: CUBIC-L^41,42^, which provided exceptional delipidation across brains of all ages (**Figure S1D**) at the cost of tissue expansion and possible damage; SHIELD^43^, which made the tissue more resilient and reduced expansion caused by CUBIC-L (**Figure S1E**); and RIMS^44^, a gentle clearing solution with adjustable refractive index (RI) and pH that also preserved EGFP. In addition, 40% urea was added to RIMS (uRIMS), which dramatically improved resolution at depth (**Figure S1C**). As highly saturated solutions (e.g. uRIMS, EasyIndex) can become turbulent due to evaporation (**Figure S1F**), a thin layer of sunflower oil was added to the surface of the chamber to prevent evaporation, markedly improving stability during imaging sessions.

**Figure 1.**
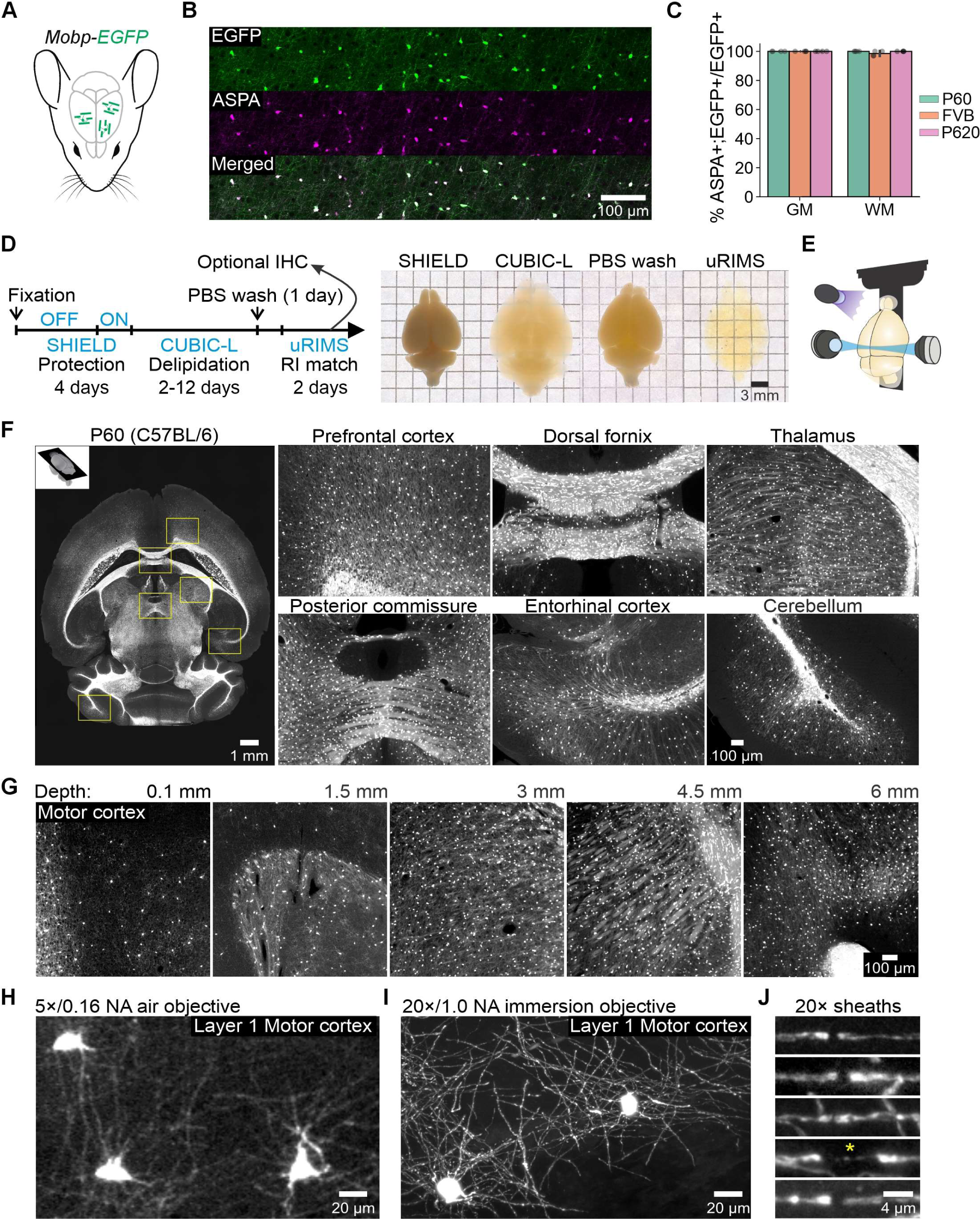
Visualization of whole brain myelin patterns. A. Oligodendrocyte somata and myelin sheaths are fluorescently labeled in *Mobp-EGFP* mice. B. Representative confocal images showing colocalization of ASPA and endogenous EGFP. C. Quantification of ASPA colocalization across mouse strain and age in gray and white matter. D. Schematic illustration of clearing pipeline (left) along with images of cleared brains at each step (right). E. Schematic of brain mounting with UV light (purple) activated glue and 3D printed sample holders. Lightsheet illumination objectives scan perpendicular to sample. F. Representative FOVs of single horizontal plane from lightsheet volume acquired with low resolution 5×/0.16 NA air objective (left). FOVs highlight the diversity of myelin patterns across brain regions (insets right). G. Representative images starting from superficial motor cortex and progressing to deeper hypothalamic regions near midline that are furthest from both illumination and detection objectives. H. Low resolution FOV of superficial cortex acquired with 5× objective. I. High resolution FOV of superficial cortex acquired with 20× objective. J. Nodes of Ranvier and paranodal bridge (asterix) visualized with 20× objective.

A minimum of two days were required to render brains optically clear using this sequential clearing pipeline, with increasing time needed to clear brains from older mice (**Figure 1D**). Samples processed and imaged in this manner (**Figure 1E**) could then be subjected to cryo-sectioning and immunocytochemistry for further analysis (**Figure S1G**). The signal-to-noise obtained in these brains with lightsheet imaging was remarkably high across all regions and depths (**Figure 1F,G; Video S1-3**), with signal loss apparent only in deeper regions of aged brains (> P620). Moreover, while myelin sheaths could be resolved with a 5× objective (**Figure 1H**), high-resolution 20× imaging was able to visualize the fine structure of individual sheaths and reliably identify nodes of Ranvier (**Figure 1I,J; Video S4**). This clearing-imaging approach was similarly effective at visualizing oligodendrocytes and myelin in the isolated spinal cord (**Figure S1H; Video S5**), highlighting its broad applicability.

### Cell detection and myelin segmentation

Oligodendrocytes account for approximately ∼20% of brain cells and are estimated to number in the millions in the adult murine brain^45^, making manual annotation impractical. Deep neural networks are particularly suited for the analysis of large fluorescence datasets, since their adaptive nature allows them to learn from differences in imaging conditions, experimental perturbations, and artifacts, to standardize analysis across datasets. However, the high density of oligodendrocytes and close spacing of their cell bodies in some brain regions presents a substantial challenge for application of many deep learning tools, such as the UNet^46^, which excels at semantic segmentation, but requires post-processing to delineate individual objects in regions of high density^47^. Therefore, we adapted a 3D Mask R-CNN algorithm^48,49^ to perform volumetric instance segmentation of individual oligodendrocyte somata. This approach provided both voxel-wise semantic segmentation and bounding box predictions, enabling the CNN to discriminate individual cells within cell clusters (**Figure 2A and S2A**). To facilitate cell detection, we applied our algorithm using an overlapping sliding window approach, ensuring that the highest resolution data was provided to the CNN and circumventing the challenge of jointly processing massive datasets, as GPU limitations prevented direct analysis of each ∼400 GB whole brain volume. All detections along window edges were then pooled together using weighted-box clustering^50^ and box splitting to remove overlap and segment individual cells of interest (**Figure 2B,C**). After analyzing whole brains, we extracted large, cropped volumes for validation and observed that the Mask R-CNN with sliding window pooling achieved a low error rate across datasets (gray matter: 0.6% error of N=1,490 detected cells; white matter: 4.3% error of N=3,113 detected cells, **Figure 2D; Video S6**). The performance of the Mask R-CNN was also superior to that of a trained UNet (**Figure S2B)** when evaluated on a set of validation patches not used for training (n=286 patches, 128 x 128 x 16 voxels XYZ, **Figure 2E**). As expected, since the UNet is designed for semantic segmentation, it achieved a higher Jaccard overlap value (UNet 0.64 ± 0.01; Mask R-CNN 0.53 ± 0.01; *p=*1.65e-9; unpaired two-tailed t-test), but was unable to accurately identify individual oligodendrocytes (Sensitivity: UNet 0.64 ± 0.02; Mask R-CNN 0.77 ± 0.02, *p=*8.1e-8; Precision UNet 0.48 ± 0.02; Mask R-CNN 0.87 ± 0.02, *p=*1.3e-49; n=286 patches, unpaired two-tailed t-test, **Figure 2F**).

**Figure 2.**
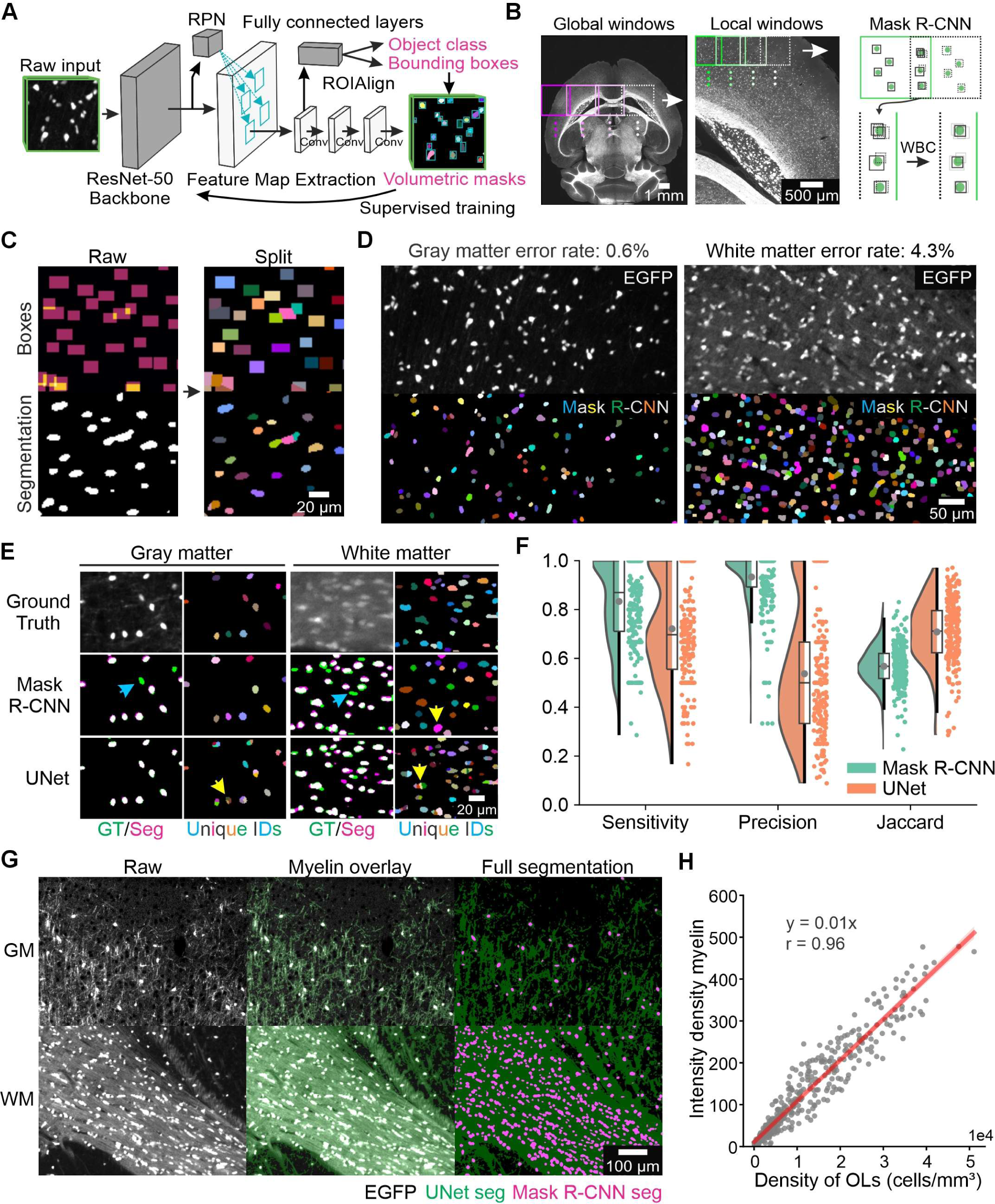
Oligodendrocyte and myelin segmentation. A. Mask R-CNN overview depicting raw input volumes (128 x 128 x 16 voxels) from which bounding boxes and semantic segmentations of individual oligodendrocytes are derived. B. Schematic of sliding window approach for analysis of BigData volumes. High resolution patches are used for analysis, and weighted box clustering (WBC) is used to accumulate detections along patch edges. C. Example of box splitting to separate overlapped regions (orange) that occur due to the rectangular shape of boxes. Once split, non-overlapping boxes are then used to mask out the binary segmentation to identify individual cells (colored randomly). D. Performance of Mask R-CNN in large, cropped volumes from gray and white matter (gray matter 550 x 450 x 1300 µm XYZ; white matter 200 x 200 x 1300 µm XYZ) using sliding window approach (gray matter: 0.6% error of n=1,490 detected cells; white matter: 4.3% error of n=3,113 detected cells). E. Comparison of Mask R-CNN and UNet on patches from validation dataset. As only a single slice is depicted for each patch, Mask R-CNN detections can appear missing on a certain slice (cyan arrows) but are not likely to be completely missed. Rather, cluster-splitting errors (yellow arrows) are the most common for both false positive and false negative detections. F. Quantification of comparison depicting sensitivity, precision, and Jaccard overlap values between detections and ground truth. G. Myelin segmentation using trained UNet. Cell somata are excluded by using previously identified cells from Mask R-CNN analysis. H. Comparison of myelin intensity density and oligodendrocyte cell density across all regions of the Allen CCF (r=0.96, *p=*8.66e-219, n=402 regions).

In *Mobp-EGFP* mice, EGFP accumulates in the cytoplasmic myelinic channels along the length of each myelin segment (inner and outer tongues) and in the paranodal loops^51^, providing a means to also estimate the density of myelin in different brain regions. To quantify brain-wide myelin densities, we trained a UNet to identify EGFP+ voxels and excluded oligodendrocyte somata previously identified by the Mask R-CNN (**Figure 2G**). Myelin intensity density was then calculated by multiplying the mean intensity for a given region by the number of EGFP+ voxels and then dividing this value by the overall volume of an area (excluding the volume occupied by oligodendrocyte somata). There was close correspondence between myelin intensity density measured through this method and absolute oligodendrocyte density (r=0.96, *p=*8.66e-219, n=402 regions, **Figure 2H**), supporting the assumption that each oligodendrocyte produces, on average, a similar amount of myelin. However, the relationship between myelin volume per region and oligodendrocyte density became non-linear in regions of high myelin density, where volume had saturated and only intensity could be used to represent increases in myelin content (r=0.88, *p=*2.9e-131, n=402 regions, **Figure S2C**). Thus, as intensity can vary substantially across tissue and between different samples (due to differences in laser settings, optics, tissue clarity, etc.), we focused on comparing oligodendrocyte density rather than myelin intensity density, as cell detection provides a more reliable means to define region specific differences.

### A computational pipeline for whole brain stitching, preprocessing and alignment to a common atlas

To enable brain-wide comparisons from terabyte-sized datasets, it is critical to establish an accurate preprocessing and registration pipeline. To assemble and register these 3D volumes to the Allen mouse brain common coordinate framework (CCF)^52^, we employed the BigStitcher platform^53^ and used affine stitching transforms to account for optical aberrations (**Figure 3**). These preprocessing steps also served to compress whole brain datasets from ∼2 TB to ∼400 GB. High resolution EGFP volumes were then accessed for deep learning analysis, while downsampled autofluorescence volumes (collected at 638 nm excitation) were extracted for registration to the Allen CCF after standard preprocessing steps (**Figure S3A**). Registration was performed using the BrainReg platform^54–56^, as diffeomorphic registration using Advanced Normalization Tools (ANTs)^57^ caused excessive warping in most cases (**Figure S3B**). A significant source of registration error was derived from mismatched autofluorescence, which was most conspicuous in white matter (**Figure S3C**). Inappropriate assignment of portions of white to gray matter at borders led to erroneous density values in neighboring gray matter areas, which was corrected using a separate mask (**Figure S3D,E,** see Methods), yielding a registration accuracy at 20 µm/pixel resolution of < 2 pixels on average (**Figure S3F**). Additional subtractions were performed to remove autofluorescent particles and residual blood present within vessels from older brains (> P620, **Figure S3I,J**, see Methods). After preprocessing, registration, and deep learning analysis, the extracted metadata was visualized as density maps and raw data was converted to N5 files that are accessible through Neuroglancer.

**Figure 3.**
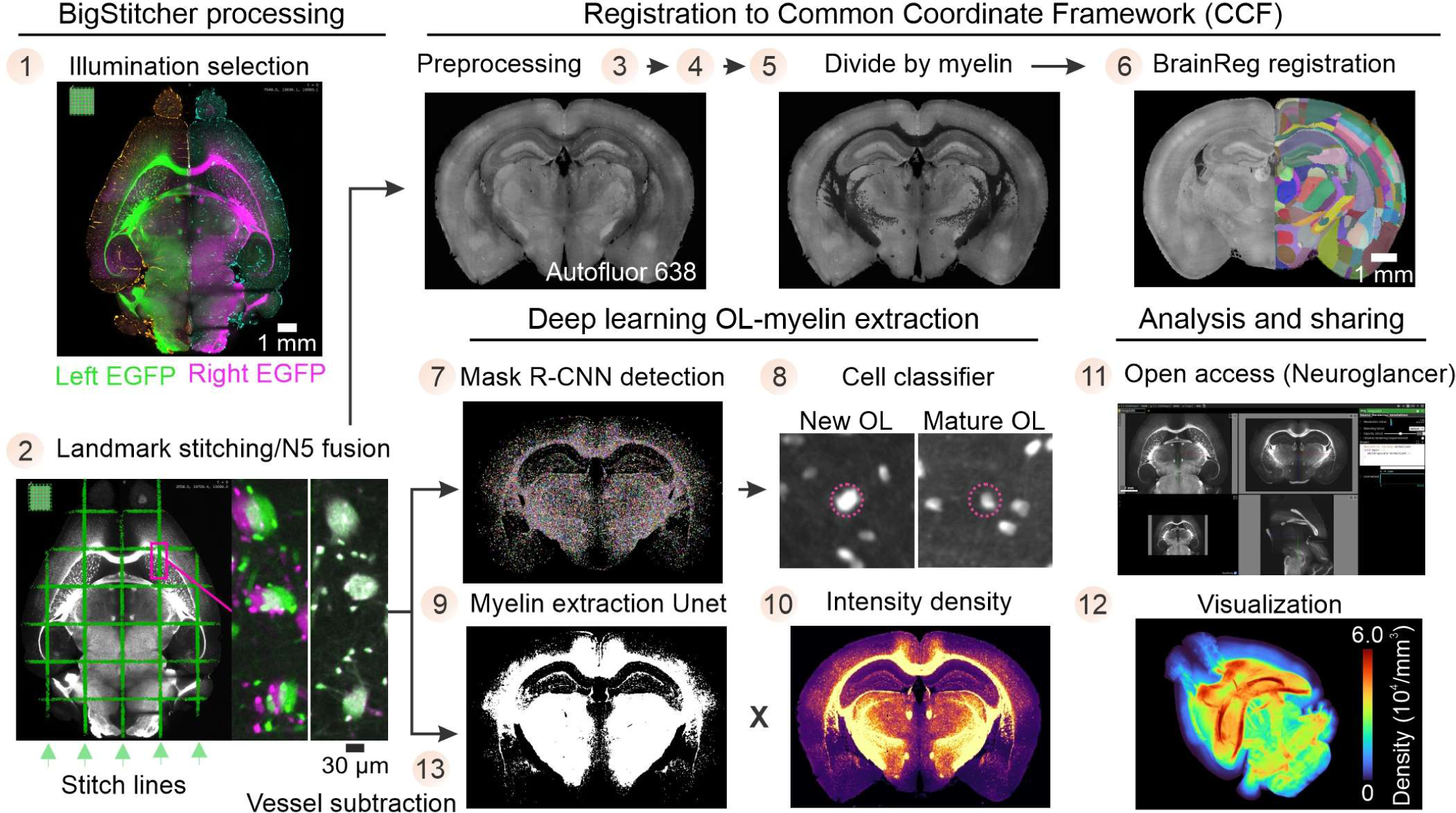
Pre-processing and registration. Diagram for computational processing pipeline of whole brain datasets. BigStitcher platform enables illumination selection and affine stitching (1-2). Extracted high resolution *Mobp-EGFP* tiles are used for oligodendrocyte and myelin segmentation (7-10, 13), while downsampled autofluorescence volumes are used for registration to the CCF using the BrainReg platform (3-6). Cross-platform compatibility between BigStitcher and Neuroglancer facilitates cloud-based data sharing of N5 files (11). Visualization of extracted counts as density maps (12).

### Oligodendrocyte distribution is conserved across hemispheres and between individuals

To define the density and distribution of oligodendrocytes within discrete brain regions, we analyzed *Mobp-EGFP* (C57BL/6) mice at postnatal day (P) 60 (**Figure 4A**), the age used to generate the Allen CCF. The density of oligodendrocytes was extremely heterogenous between brain regions at this age, ranging across four orders of magnitude (**Figure 4B** and **S4A**). CCF delineated functional boundaries often matched transitions in oligodendrocyte densities between neighboring regions, such as a dearth of oligodendrocytes in deep layers of the temporal association (TEa), ectorhinal (ECT), and perirhinal (PERI) cortices (**Figure 4C**). These patterns were remarkably conserved across hemispheres (r=0.99, *p=*2.7e-413, n=417 regions, **Figure 4D**), and between individuals, with the average deviation from the mean of ∼10% (defined as coefficient of variation, CV) for most regions (**Figure 4E**).

**Figure 4.**
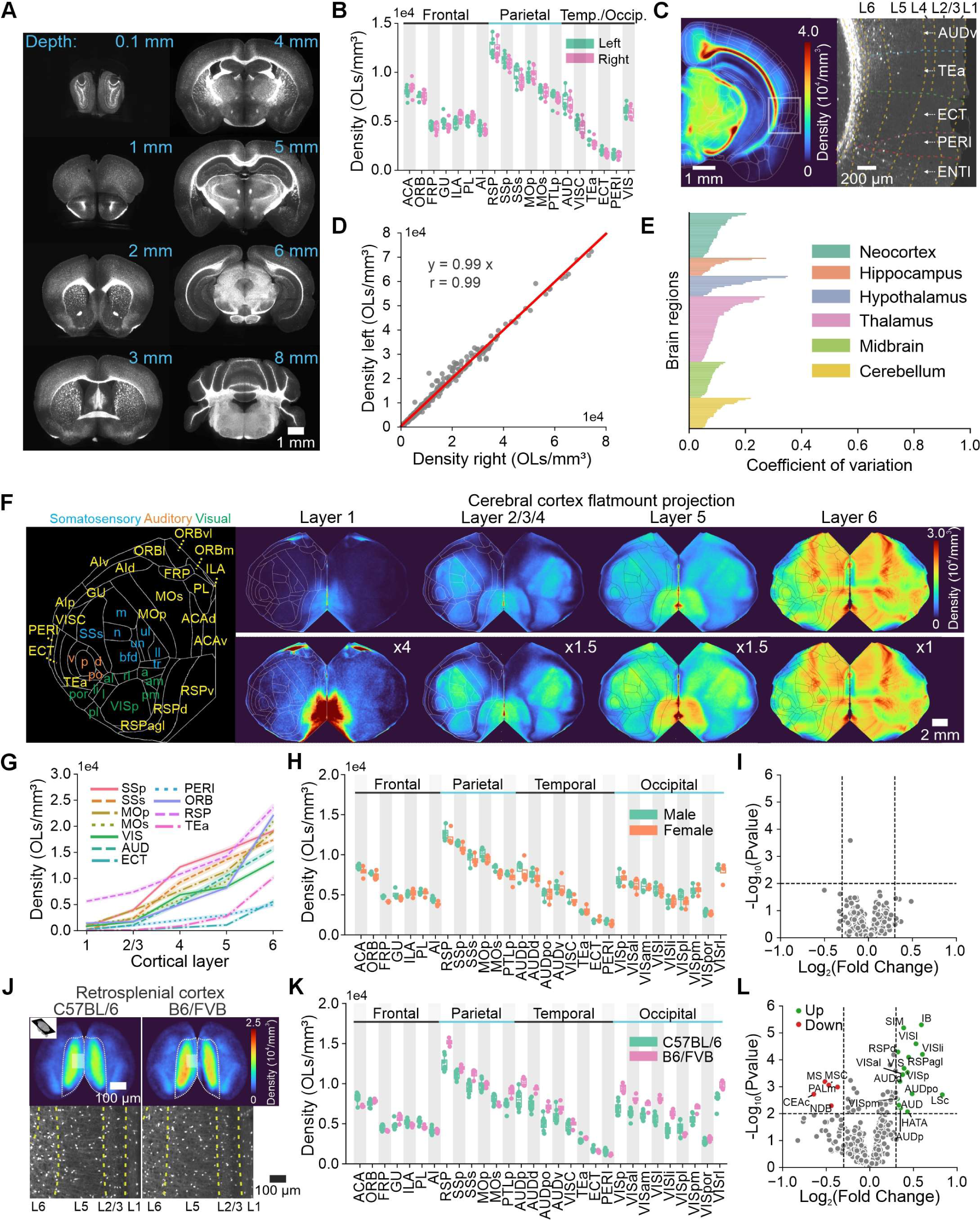
Oligodendrocyte density across hemispheres, individuals, sex, and strain. A. Sequential coronal sections across a whole *Mobp-EGFP* brain highlighting bilateral symmetry. B. Oligodendrocyte density across all regions of neocortex (ACA–anterior cingulate, ORB–orbital, FRP–frontal pole, GU–gustatory, ILA–infralimbic, PL–prelimbic, AI–agranular insular, RSP– retrosplenial, SSp–primary somatosensory, SSs–supplemental somatosensory, MOp–primary motor, MOs–secondary motor, PTLp–posterior parietal association, AUD–auditory, VISC– visceral, TEa – temporal association, ECT – ectorhinal, PERI – perirhinal, VIS – visual, n=8). C. Cross-section of average density map and inset showing low density of oligodendrocytes in L6 of temporal association (TEa), ectorhinal (ECT), and perirhinal (PERI) regions. D. Strong correlation between myelin density across left and right hemispheres (r=0.99, *p=*2.7e-413, n=417 regions averaged across 8 mice). E. Coefficient of variation (CV) across brain regions to show deviation from mean (CV neocortex: 0.1 ± 0.01; hippocampus: 0.11 ± 0.02; hypothalamus: 0.17 ± 0.03; thalamus: 0.11 ± 0.01; midbrain: 0.08 ± 0.004; cerebellum: 0.1 ± 0.01, n=8 mice). F. Cortical flatmount projections of average cell density maps for P60 brain (n=8) across each layer of the cortex. Cell density is scaled as indicated (bottom row) to better visualize differences across more superficial layers that are less myelinated. Additional subregions of somatosensory cortex (SS) (m–mouth, n–nose, bfd–barrel field, un–unassigned, ul–upper limb, ll–lower limb, tr–trunk), orbital cortex (ORB) (vl–ventrolateral, m–medial, l–lateral), and additional acronyms (d–dorsal, v–ventral, agl–lateral agranular, p–primary, po–posterior, al– anterolateral, am–anteromedial, li–laterointermediate, pl–posterolateral, pm–posteromedial, por–postrhinal, rl–rostrolateral). G. Oligodendrocyte density across cortical layers and regions (n=8). H. Comparison of cortical oligodendrocyte density across all regions of neocortex between sexes. I. Volcano plot of all brain regions showing no significant differences across sex (p-value threshold = 0.01; Log_2_ fold change threshold = 0.3, n=4 males and n=4 females). J. Retrosplenial cortex density maps from C57BL/6 and mixed background B6/FVB brains. K. Comparison of oligodendrocyte densities across all regions of neocortex between C57BL/6 and mixed background B6/FVB brains. L. Volcano plot of all brain regions showing statistically significant differences in cell density when comparing B6/FVB mice to C57BL/6 animals. “Up” indicates a higher density of oligodendrocytes in B6/FVB mice (p-value threshold = 0.01; Log_2_ fold change threshold = 0.3, n=8 B6 mice and n=4 mixed B6/FVB mice).

The neocortex is among the last regions to be myelinated during development^58^ and exhibits a distinct laminar structure, allowing further examination of oligodendrocyte distribution within well-defined neuronal subnetworks. To facilitate layer-specific comparisons between cortical regions, we generated digital “flatmount” projections, where all cortical regions are represented in a two-dimensional map for each layer (**Figure 4F and S4B**). These projections revealed previously reported trends, such as the progressive increase in oligodendrocyte density from layer (L)1 to L6 (**Figure 4F,G and S4C**); however, individual regions exhibited marked differences in oligodendrocyte abundance. For instance, L2/3/4 regions that receive direct sensory input (L4-primary somatosensory (SSp), L4-primary visual (VISp), L4-primary auditory (AUDp)) had up to three times more oligodendrocytes than neighboring areas that often lacked a functionally distinct L4, such as primary motor cortex (MOp) (cell density per mm^3^ in L2/3-MOp: 3,891 ± 212 compared to L4-SSp: 12,198 ± 209, *p=*1.13e-13; L4-VISp: 7,890 ± 366, *p=*1.85e-7; L4-AUDp: 6,272 ± 588, *p=*0.002; n=8 mice for all comparisons, unpaired two-tailed t-test, **Figure 4F,G**). Moreover, while oligodendrocyte density was higher in L1 of the retrosplenial cortex (RSP) than in all other cortical regions, this relationship was not consistent across deeper layers, as L6-orbital (ORB), L6-primary somatosensory (SSp), and L6-secondary motor (MOs) cortices all had similar oligodendrocyte densities to L6-RSP cortex (cell density per mm^3^ in L6-RSP: 23,608 ± 596 compared to L6-ORB: 22,053 ± 409, *p=*0.05; L6-SSp: 19,110 ± 265, *p=*0.002; L6-MOs: 20,952 ± 385; n=8 mice, unpaired two-tailed t-test, **Figure 4G**). These findings highlight the extreme regional diversity in oligodendrocytes across functionally distinct cortical layers and regions, contrasting with the near-uniform density of OPCs across the cortical mantle^5,59^.

### Oligodendrocyte distributions are conserved between sexes but vary between strains

Sexual dimorphisms in myelin content have been reported in human patients based on diffusion tensor imaging (DTI), though often contradictory or inconclusive^9^, and are associated with differential rates of prevalence for myelin-associated diseases like multiple sclerosis (MS)^60^. Nevertheless, there was remarkable concordance in global oligodendrocyte densities between sexes in the P60 C57BL/6 murine brain (**Figure 4H**), and even regions that exhibit sexual dimorphism (e.g. amygdala, hypothalamus^61^) had similar oligodendrocyte densities (cell density per mm^3^ in cortical amygdalar area (COA): male 1,720 ± 256, female 1,568 ± 31, *p=*0.58, n=4 per group, unpaired two-tailed t-test; Hypothalamus (HY): male 19,241 ± 1,226; female 17,697 ± 1,410, *p=*0.44, n=4 mice per group, unpaired two-tailed t-test), as shown by the tight clustering of regional densities in volcano plots (**Figure 4I**). Moreover, as male mice were larger than females (weights (g): male 22.4 ± 1.7; female 17.9 ± 0.83, n=4 per group), mouse size also did not appear to correlate with myelin complexity.

Genetic differences between mouse strains are associated with differences in metabolism^62^, immune responses^63^, and neurological features, such as the response to anesthesia^64^, injury and neurodegenerative diseases^65^. Differences between C57BL/6 and FVB/NJ mice are particularly well documented, with significant variability in sensorimotor performance, learning, and memory^66^. Additional mutations also cause gradual hearing loss in C57BL/6 animals^67^ and retinal degeneration in FVB/NJ mice^68^. To explore whether regional differences in oligodendrocyte density vary with genetic background, we crossed *Mobp-EGFP* C57BL/6 mice with wild type FVB/NJ mice to generate mixed background (B6/FVB) *Mobp-EGFP* animals. Brains were then collected at P60 and subjected to clearing, imaging, and quantification (**Figure S4D**). Statistically significant differences in oligodendrocyte densities between retrosplenial (RSP, cell density per mm^3^ in B6 mice: 12,374 ± 334, n=8; FVB mice: 15,176 ± 229, n=4, *p=*0.0003), primary visual (VISp, cell density per mm^3^ in B6 mice: 6,994 ± 279, n=8; FVB mice: 9,426 ± 293, n=4, *p=*0.0003), and primary auditory regions (AUDp, cell density per mm^3^ in B6 mice: 8,015 ± 425, n=8; FVB mice: 10,235 ± 270, n=4, *p=*0.0006) were detected after just one generation of breeding (**Figure 4J,K**), visible in the greater spread of points (corresponding to brain regions) in the volcano plot (**Figure 4L**). Notably, these variations presented as relatively uniform increases across whole regions in the density map (**Figure 4J**), rather than local shifts in density gradients within a region, suggesting that they may reflect differences in rates of development.

### Oligodendrocyte addition across the lifespan preserves regional differences in density

Oligodendrocytes are produced throughout life from a resident population of progenitor cells (OPCs), resulting in a gradual increase in oligodendrocyte and myelin density with age^10–12^. Although previous studies have documented age-dependent changes in the density of oligodendrocytes and myelin in individual brain regions^10^, there have been no brain-wide quantitative developmental studies of oligodendrocytes. To determine how the extended period of oligodendrogenesis influences regional differences in oligodendrocyte densities observed at P60, we selected mice of three additional ages (P240, P620, P850) for detailed analysis, covering the lifespan of most mice. Ambiguous regions were excluded from analysis at P850, where tissue and image quality compromised assessment within deeper brain regions (**Table S1**). Heat map projections revealed that there was a general increase in oligodendrocyte density with age across broadly defined brain regions (**Figure 5A**), with the highest density change in fiber tracts and the highest fold increases occurring in the neocortex and hippocampus (**Figure 5B**). Notably, the higher normalized fold change in neocortex was not simply a reflection of having fewer cells at baseline, as the cerebellar cortex also had a low density of oligodendrocytes at P60, but the normalized fold change remained low across all ages (**Figure 5B**). Thus, the high fold change in neocortex and hippocampus reflected their considerably more permissive environments for oligodendrogenesis.

**Figure 5.**
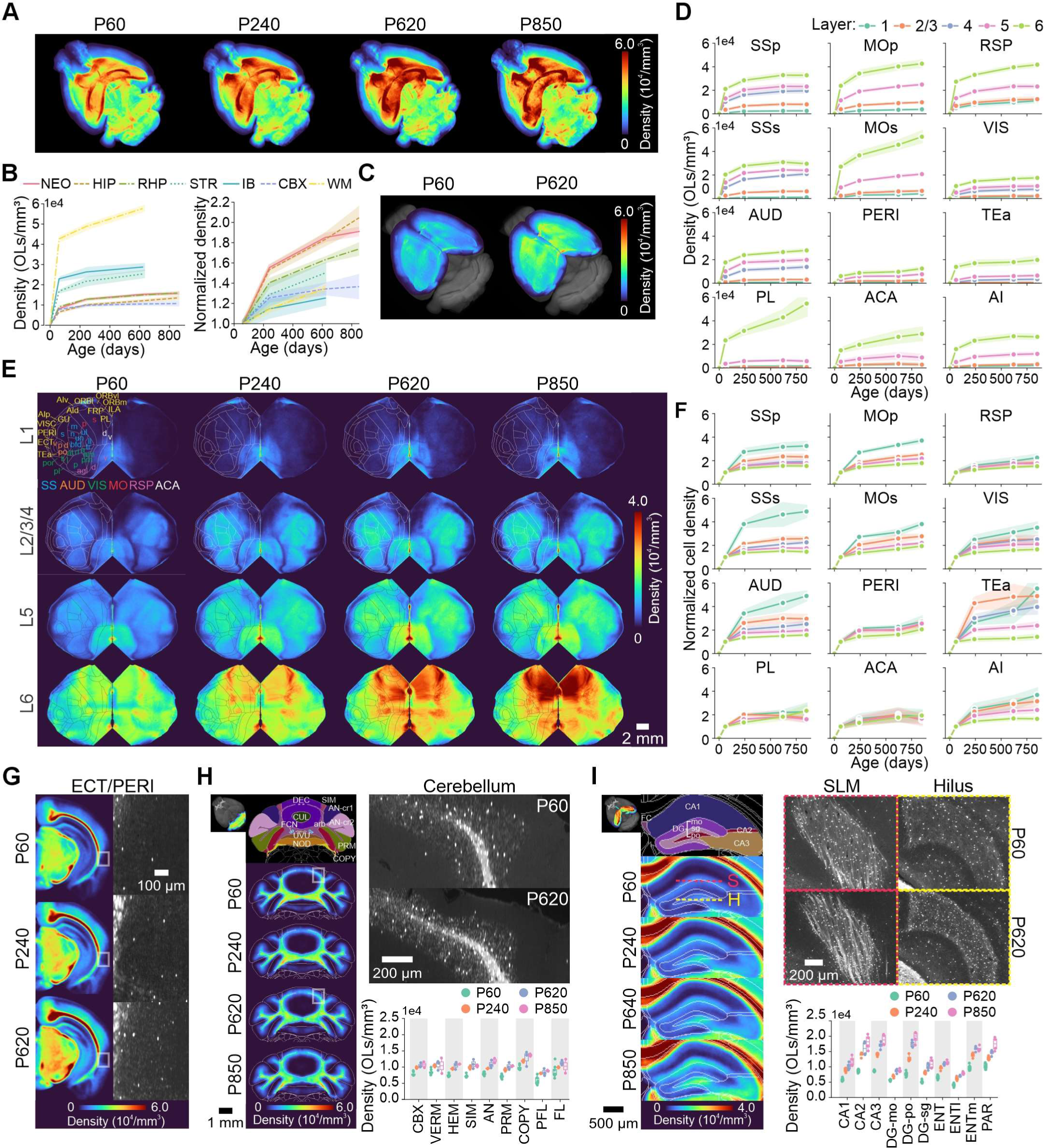
Aging myelin landscape. A. Average cell density maps at each timepoint across the murine lifespan. B. Global comparisons of oligodendrocyte density across broadly defined regions. Absolute density (left) (rank order change in mean cell density from P60 to P620: white matter (WM) +14,965; striatum (STR) +8,456; neocortex (NEO) +7,029; interbrain (IB) +5,802; retrohippocampal area (RHP) +5,787; hippocampus (HIP) +5,491; cerebellar cortex (CBX) +2,690) and normalized density (right) (rank order mean fold change from P60 to P620: neocortex 1.85; hippocampus 1.83; retrohippocampal area 1.63; striatum 1.5; white matter 1.35; cerebellar cortex 1.34; interbrain 1.25). C. Density map over time focused on neocortex overlaid on Allen CCF autofluorescence (gray). D. Comparison of changes in oligodendrocyte density across select cortical regions and layers with age (SSp–primary somatosensory, MOp–primary motor, RSP–retrosplenial, SSs–supplemental somatosensory, MOs–secondary motor, VIS–visual, AUD–auditory, PERI–perirhinal, TEa– temporal association, PL–prelimbic, ACA–anterior cingulate, AI–agranular insular, n=8 for P60 and n=4 for all other ages). E. Cortical flatmounts of average cell density maps across layers with age (n=8 for P60 and n=4 for all other ages). F. Comparison of normalized fold change in oligodendrocyte density with age. G. Coronal sections of average cell density maps and raw data (insets) of ectorhinal/perirhinal areas across aging. H. Coronal sections of cerebellar cortex and raw data (insets) across aging, with associated density comparisons across select sub-regions of cerebellar cortex (CBX–cerebellar cortex, VERM–vermal regions, HEM–hemispheric regions, SIM–simple lobule, AN–ansiform lobule, PRM–paramedian lobule, COPY–copula pyramidis, PFL–paraflocculus, FL–flocculus). I. Coronal sections of hippocampus and raw data (insets) of hilus (H) and stratum lacunosum-moleculare (S) across aging. While the Allen CCF at 20 µm/px resolution does not provide the delineated borders of the stratum lacunosum-moleculare, it is easily identifiable by its location adjacent to the dentate gyrus and the myelinated fiber bundles within. Associated plots of density comparisons across select sub-regions of hippocampus (CA1, CA2, CA3, Dentate gyrus (DG)-mo–molecular layer, DG-po–polymorph layer, DG-sg–granule cell layer, ENT–entorhinal area, ENTl–lateral entorhinal, ENTm–medial entorhinal, PAR–parasubiculum).

Overall, the neocortex exhibited a rapid increase in density at earlier ages (P0–P240), plateauing by P620, with no noticeable inflections to indicate multi-phasic trends or significant cell loss (**Figure 5C-F and S5A,B**). However, when examining subregions more closely, we observed layer-specific changes that deviated from these general trends. For instance, oligodendrocyte density continued to increase rapidly in L6-secondary motor (MOs) and L6-prelimbic (PL) areas of the prefrontal cortex into old age (**Figure 5D,E and S5C**). Moreover, the fold change in oligodendrocyte density was not the same across cortical layers; the highest initial changes were observed in L1 and decreased sequentially from L1 to L6 (**Figure S5D**). Conversely, while the fold change was highest in L1, the absolute change in cell density was highest in L4 and L6 with age (**Figure S5E**).

Despite these layer-specific deviations, regional differences between cortical areas were remarkably preserved over the lifespan: there was no correlation between cell density at P60 and fold change at P620 (r=-0.17, *p=*0.35, n=34 regions, **Figure S5F**), as the average fold change was similar for most regions. These results indicated that the neocortex maintains the overall, differential distribution of oligodendrocytes as the brain ages, such that regions with low density remain low and do not “catch up” to highly myelinated areas. Low density regions, like the perirhinal cortex, increased proportionally with age, but remained largely sparsely populated by oligodendrocytes (**Figure 5G**). Moreover, the relative difference in oligodendrocyte density between primary and supplemental areas (SSp vs. SSs and MOp vs. MOs) was maintained across the lifespan (**Figure S5G**, ratio of cell density in primary vs. supplemental areas; SSp-SSs ratio at P60: 1.17 and ratio at P620: 1.18; MOp-Mos ratio at P60: 1.22 and ratio at P620: 1.22). However, when comparing the neocortex to non-cortical regions, the rate of change was more heterogenous. For instance, while the cerebellar cortex had a similar density of oligodendrocytes at P60 to the neocortex, the relative fold change in the cerebellar cortex was much smaller than in the neocortex with aging (**Figure 5B,H**), highlighting that dramatic regional differences in oligodendrogenesis persist throughout the lifespan.

### Regional heterogeneity in oligodendrocyte distribution within the hippocampus

The hippocampus plays a critical role in memory consolidation and spatial learning, and is susceptible to hyperactivity in epilepsy^69^, but the developmental organization of oligodendrocytes within this structure and the targets of myelination have not been comprehensively defined^70^. Notably, the principal mossy fiber and Schaffer collateral projections are not myelinated^71,72^. Like the neocortex, the hippocampus experienced a large fold change in density of oligodendrocytes with age (**Figure 5B**). The regional specificity of these changes was most striking in the dentate gyrus, as the hilus (DG-po) contained significantly more oligodendrocytes than neighboring molecular (DG-mo) and granule cell (DG-sg) layers across all ages and experienced a more dramatic increase over the lifespan (All comparisons with DG-po; DG-mo at P60 (*p=*4.9e-6), P850 (*p=*3.5e-5); DG-sg at P60 (*p=*1.2e-6), P620 (*p=*6.6e-5), n=8 for P60 and n=4 for all other ages, unpaired two-tailed t-test, **Figure 5I**). Unlike the distribution of myelin sheaths within the stratum lacunosum-moleculare (SLM) of area CA1 (not defined in 20 µm Allen CCF), which was arranged into thick bundles, oligodendrocyte somata and myelin in the hilus were distributed more evenly (**Figure 5I**), possibly corresponding to myelination of interneuron axons. A striking gradient in oligodendrocyte density was also visible in the entorhinal cortex, with the medial region (ENTm) containing more than twice the density of oligodendrocytes as the lateral region (ENTl) (cell density per mm^3^ at P60 for ENTl: 4,045 ± 178 and ENTm: 9,648 ± 292, n=8, *p=*1.6e-10, unpaired two-tailed t-test, **Figure 5I**). Together, these findings highlight the extreme regional diversity in oligodendrocytes and extended time course of myelin patterning within these temporal lobe structures.

### Cuprizone injury and recovery dynamics vary markedly across the brain

Understanding the mechanisms of oligodendrocyte injury and recovery between different brain regions is critical to develop new treatments for demyelinating disorders such as MS, in which lesions manifest across the brain. To explore the sensitivity of oligodendrocytes to injury and assess brain-wide myelin regeneration, we used our mapping pipeline to study the loss and recovery of oligodendrocytes after exposure to the oligotoxin cuprizone. *Mobp-EGFP* mice (P60) were exposed to 0.2% cuprizone for six weeks to ablate oligodendrocytes and then sacrificed or allowed to recover for an additional three weeks to visualize the initial stages of oligodendrocyte regeneration (**Figure S6A**). After six weeks of cuprizone, all areas of the brain experienced a substantial loss of oligodendrocytes (**Figure 6A**), with the largest absolute loss in the interbrain (thalamus and hypothalamus) and the largest normalized fold change in the neocortex and hippocampus (**Figure 6B**). Similar to the developmental progression of oligodendrocyte integration, this large fold change in neocortex was not simply a consequence of the low cell density at baseline, as the cerebellar cortex had an equally low density of oligodendrocytes at P60, but experienced minimal fold change loss following cuprizone (**Figure 6B**). Recovery dynamics were also strikingly variable between brain regions, with the cerebellum and interbrain regions exhibiting substantially more complete recovery than more depleted regions like the neocortex and hippocampus (**Figure 6B**), indicating that the degree of injury does not indicate the rate of recovery. These disparities in global injury-recovery responses suggest that local factors, such as neural activity or inflammation^4,73^, may greatly influence the survival and regeneration of oligodendrocytes.

**Figure 6.**
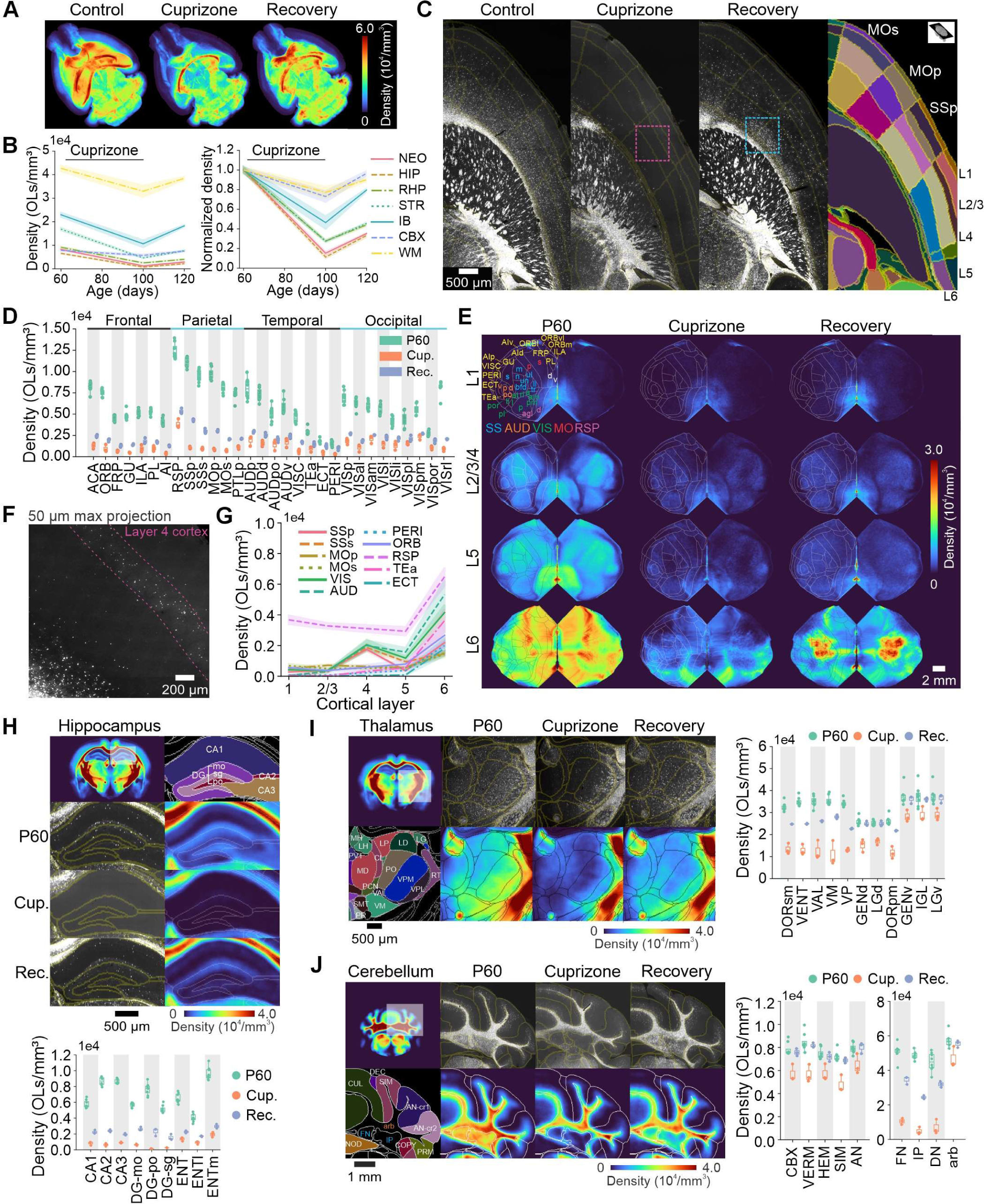
Cuprizone injury-recovery dynamics are asynchronous. A. Average density maps comparing oligodendrocyte density immediately after six weeks of cuprizone injury and after three weeks of recovery (n=8 for control and n=3 mice for cuprizone and recovery conditions). B. Global comparison of oligodendrocyte density across broadly defined regions immediately after cuprizone and during recovery. Absolute density (left) (rank order change in mean cell density per mm^3^ after six weeks of Cuprizone: interbrain (IB) -12,307; striatum (STR) -12,230; white matter (WM) -9,711; neocortex (NEO) -6,997; retrohippocampal area (RHP) -6,626; hippocampus (HIP) -5,847; cerebellar cortex (CBX) -2,120) and normalized density (right) (rank order mean normalized density change after six weeks of cuprizone: -0.89 hippocampus; -0.85 neocortex; -0.73 striatum; -0.72 retrohippocampal area; -0.54 interbrain; -0.27 cerebellar cortex; -0.23 white matter). C. Horizontal cross-section of the neocortex after injury and during recovery with CCF boundaries (right). Magenta box highlights layer 4 inset during injury (Figure 6F). Cyan box highlights layer 6 inset during recovery (Figure S6C). D. Side-by-side comparisons of oligodendrocyte density after injury and during recovery for all regions of neocortex (ACA–anterior cingulate, ORB–orbital, FRP–frontal pole, GU–gustatory, ILA–infralimbic, PL–prelimbic, AI–agranular insular, RSP–retrosplenial, SSp–primary somatosensory, SSs–supplemental somatosensory, MOp–primary motor, MOs–secondary motor, PTLp–posterior parietal association, AUD–auditory, VISC–visceral, TEa–temporal association, ECT–ectorhinal, PERI–perirhinal, VIS–visual). Additional auditory sub-regions (AUDp–primary auditory, d–dorsal, po–posterior, v–ventral) and visual sub-regions (VISp– primary visual, al–anterolateral, am–anteromedial, l–lateral, li–laterointermediate, pl– posterolateral, pm–posteromedial, por–postrhinal, rl–rostrolateral). E. Cortical flatmount projections of average cell density maps after injury and during recovery. F. Inset showing surviving oligodendrocytes that form distinct line in layer 4 sensory regions. G. Density of oligodendrocytes by layer after injury (pooled from n=3 mice). H. Coronal section of average density maps and raw *Mobp-EGFP* fluorescence for hippocampus, with associated plot of density comparisons for select regions (CA1, CA2, CA3, Dentate gyrus (DG)-mo–molecular layer, DG-po–polymorph layer, DG-sg–granule cell layer, ENT–entorhinal area, ENTl–lateral entorhinal, ENTm–medial entorhinal, PAR–parasubiculum). I. Coronal section of thalamus density map and raw fluorescence with associated plot of density comparisons for select regions of sensory-motor cortex related thalamus (DORsm): VENT– ventral group of dorsal thalamus, VAL–ventral anterior-lateral complex, VM–ventral medial nucleus, VP–ventral posterior complex, GENd–geniculate group of dorsal thalamus, LGd–dorsal lateral geniculate complex. Also shown are select regions of polymodal association cortex related thalamus (DORpm): GENv–genicular group of ventral thalamus, IGL–intergeniculate leaflet, LGv–ventral lateral geniculate complex). J. Coronal section of cerebellum density map and raw fluorescence with associated plot of density comparisons for select regions (CBX–cerebellar cortex, VERM–vermal, HEM–hemispheric, SIM–simple lobule, AN–ansiform lobule) including deep cerebellar nuclei (FN–fastigial nucleus, IP–interposed nucleus, DN–dentate nucleus—in addition to cerebellar white matter (arbor vitae, arb).

Cuprizone induced demyelination has been studied extensively within the cerebral cortex^70,74,75^, but oligodendrocyte susceptibility to this toxin and their regeneration after loss have not been systematically compared between cortical regions and across distinct cortical layers. Whole cortex analysis revealed that the extent of oligodendrocyte death varied across cortical regions (**Figure 6C,D**), with a pattern highly concordant between individuals. For instance, retrosplenial cortex (RSP) was consistently less susceptible to cuprizone compared to similarly myelinated regions, such as primary somatosensory cortex (SSp) (cell density per mm^3^ after cuprizone injury: RSP: 3,838 ± 305 and SSp: 1,047 ± 86, n=3, *p=*9.2e-4, unpaired two-tailed t-test, **Figure 6D**), but experienced slower recovery (recovery fold change after injury: RSP: 1.39 ± 0.05 and SSp: 4.13 ± 0.08, n=3, *p=*9.0e-6, unpaired two-tailed t-test, **Figure 6D**). Moreover, the pattern of loss between cortical layers correlated with functionally distinct regions (**Figure 6E; Figure S6B**). In particular, at the end of cuprizone exposure, L4 sensory areas (SSp, SSs, VIS, and AUD) had a higher density of surviving oligodendrocytes than L1, L2/3, or L5 (**Figure 6F,G**).

The ability of different cortical regions to regenerate oligodendrocytes after three weeks of recovery was also heterogenous, with L6-SSp displaying the highest rates of oligodendrogenesis, specifically in the mouth, nose and barrel field areas (**Figure 6E; Figure S6B**). While our data recapitulated previous findings that L1 recovers proportionally the most *in vivo*, with sequentially less proportional recovery in deeper layers^4,76^, L6-SSp recovery went against this trend, and was higher than all other non-superficial SSp layers (**Figure S6F**). Notably, myelin formed by the newly generated oligodendrocytes in L6-SSp was primarily associated with laterally oriented fiber tracts that ran parallel to white matter (**Figure S6C**). Finally, while the band of surviving oligodendrocytes in L4 was apparent immediately at the end of cuprizone exposure (**Figure 6F**), both L4 and L5 consistently exhibited the lowest rates of recovery (**Figure S6E,F**) further highlighting the disconnect between oligodendrocyte loss and regeneration.

Regional differences in injury and recovery were also evident outside the cortex. The hippocampus exhibited the most significant decrease in oligodendrocyte density after cuprizone, with nearly complete ablation of all oligodendrocytes (**Figure 6H; Figure S6G**). The thalamus also displayed prominent oligodendrocyte loss but experienced more substantial recovery than the cortex (**Figure 6I; Figure S6H**). Neighboring thalamic nuclei corresponding to functionally distinct regions also displayed differential rates of injury and recovery. For instance, adjacent structures, such as the ventral medial nucleus (VM) and the ventral posterior complex (VP), had similar cell densities at baseline and injury, but VM experienced significantly more recovery (cell density per mm^3^ during recovery for VP: 22,646 ± 153 and VM: 28,018 ± 517, n=3, *p=*0.0006, unpaired two-tailed t-test, **Figure 6I**). The cerebellar cortex was a notable exception to the rest of the brain, as oligodendrocyte loss was minimal, and recovery was nearly complete (**Figure 6J; Figure S6I**). However, oligodendrocyte loss within deep cerebellar nuclei—fastigial nucleus (FN), interposed nucleus (IP), dentate nucleus (DN)—was much more prominent, with a near full ablation followed by partial recovery (**Figure 6J).** Remarkably, injury to deep cerebellar nuclei was tightly restricted spatially, as surrounding white matter regions (the arbor vitae, arb) were minimally injured (**Figure 6J**). Together, these results show how the diversity in myelin loss and recovery closely aligns to functionally distinct brain regions and sub-circuits, suggesting a prominent role for neural control of both oligodendrocyte state and oligodendrogenesis.

### Brain-wide identification of newly formed oligodendrocytes

Oligodendrocytes undergo profound morphological changes after their initial generation. As identified through *in vivo* time lapse imaging in *Mobp-EGFP* animals^77^, oligodendrocytes exhibit a gradual decrease in soma size over several weeks until they reach their mature state (**Figure 7A**). This soma volume reduction is also accompanied by a decline in EGFP fluorescence in the soma and processes^77^, reflecting both the lower cytosolic volume and lower *Mobp* promoter activity. These highly stereotyped changes provide a means to selectively identify newly generated oligodendrocytes within the vast population of mature oligodendrocytes in the brain. To automatically identify sites of active oligodendrogenesis brain-wide, we applied these principles to train a Vision Transformer (ViT) to classify oligodendrocytes as newly formed or mature. The network was provided with cropped volumes centered around the soma of candidate cells from whole brain lightsheet datasets to provide local context for decision-making (**Figure 7B,C**). Local context was critical for accurate classification, as optical aberrations warped the edges of image tiles, such that cells along stitch lines appeared larger than those at the center of an image tile, resulting in false positives. ViTs are well suited for this classification task, as the patch-based self-attention architecture allows the network to learn about spatial relationships from the provided local context^39^. This approach was able to achieve an accuracy of 84% on a validation dataset of 1,712 cells withheld from training (**Figure 7D**). To verify if the ViT was identifying large cells, we then compared the size of each ViT-identified newly formed oligodendrocyte to eight nearest-neighbor oligodendrocytes, and found that 99.9% of all ViT-identified cells were larger than the average size of their neighbors (mean volume µm^3^: ViT-classified cells 1,662 ± 3.2; nearest-neighbors 722 ± 1.4, n=25,515 cells, *p=*3.4e-9853 unpaired two-tailed t-test, **Figure 7E**).

**Figure 7.**
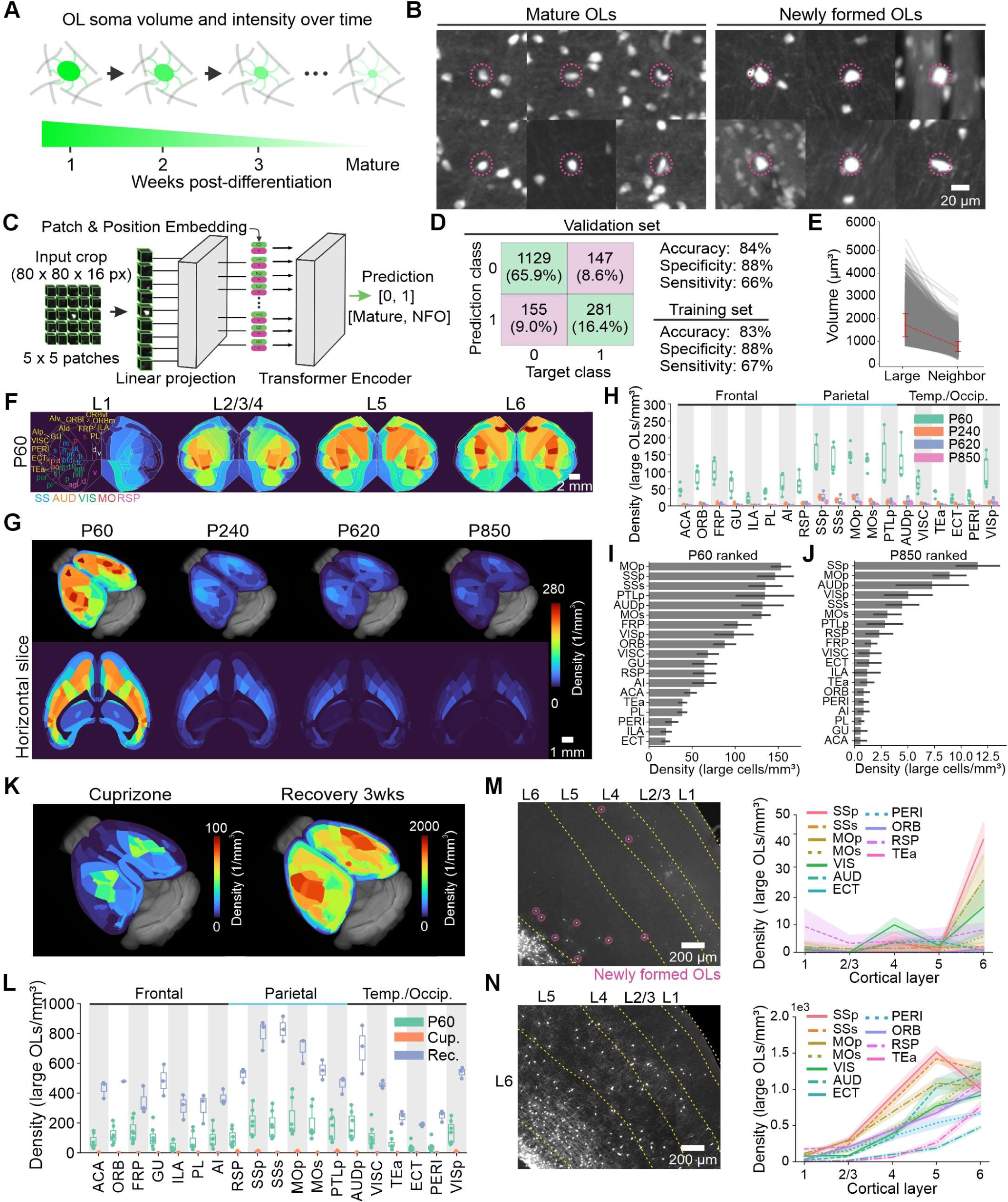
Mapping oligodendrogenesis using oligodendrocyte morphology. A. Schematic of concerted morphological changes post-differentiation. Newly formed oligodendrocyte somata are significantly larger and brighter than mature oligodendrocyte somata in *Mobp-EGFP* mice. B. Examples of mature and newly formed oligodendrocytes. C. Schematic of ViT model for classifying oligodendrocytes as mature or newly formed (NFO). D. Performance of ViT model on validation and training sets. E. Comparison of soma volume between identified large oligodendrocytes and eight nearest-neighbor cells (mean volume µm^3^: ViT-classified cells 1,662 ± 3.2; nearest-neighbors 722 ± 1.4, n=25,515 cells, *p=*3.4e-9853 unpaired two-tailed t-test). F. Region-based density maps of newly formed oligodendrocytes across cortical layers at P60. G. Region-based density maps depicting changes in density of newly formed oligodendrocytes with age. H. Density of large oligodendrocytes across all regions of neocortex at each aging timepoint ACA– anterior cingulate area, ORB–orbital area, FRP–frontal pole, GU–gustatory areas, ILA– infralimbic area, PL–prelimbic area, AI–agranular insular area, RSP–retrosplenial cortex, SSp– primary somatosensory area, SSs–supplemental somatosensory area, MOp–primary motor area, MOs–secondary motor area, PTLp–posterior parietal association areas, AUDp–primary auditory area, VISC–visceral area, TEa–temporal association area, ECT–ectorhinal area, PERI– perirhinal area, VISp–primary visual area, n=8 for P60 and n=4 mice for all other timepoints). I. Ranked list of regions with highest density of large oligodendrocytes at P60. J. Ranked list of regions with highest density of large oligodendrocytes at P850. K. Region-based density maps of large oligodendrocytes after cuprizone injury and during recovery. L. Density of large oligodendrocytes across all regions of neocortex from baseline to recovery (n=8 for P60 and n=3 for cuprizone and recovery). M. Inset of somatosensory cortex after injury, depicting newly formed oligodendrocytes (magenta circles) in L6 and L4, along with associated quantification of large oligodendrocyte density across all cortical layers and regions (right) immediately after injury. N. Inset of somatosensory cortex during recovery and associated quantification of large oligodendrocyte density across all cortical layers and regions (right).

As the density of newly formed oligodendrocytes was low and their distribution wide, we created region-based density maps to highlight differences in oligodendrogenesis between functionally distinct brain regions (**Figure 7F,G**). Analysis was restricted to the neocortex and hippocampus, as the density of oligodendrocytes was too high in white matter areas for the ViT to reliably distinguish between individual large cells and cell clusters. Applying this analysis to brains collected across the lifespan revealed the expected, marked decline in oligodendrogenesis from P60 to P850 (**Figure 7G,H**), although new oligodendrocyte production was detected in mice over two years old. This decline in oligodendrogenesis matched the aging-dependent plateau in oligodendrocyte density (**Figure 5A**), and is consistent with *in vivo* imaging studies showing that successful integration of newly formed oligodendrocytes declines with age^10^. Rank ordering of brain regions by the density of newly formed oligodendrocytes revealed high variance in oligodendrocyte production between areas, with primary sensory and motor regions maintaining the highest rates of oligodendrogenesis across the lifespan (**Figure. 7I,J**).

To identify newly regenerated oligodendrocytes in the context of injury, we applied the ViT to cuprizone and recovery datasets. As expected, the density of newly formed oligodendrocytes was substantially higher during recovery than at baseline (**Figure 7K,L**). Notably, this approach made it possible to distinguish regions with surviving oligodendrocytes from regions with high rates of oligodendrogenesis, despite having a similar overall number of oligodendrocytes. For example, although a high density of oligodendrocytes was detected in both L4 and L6 at the end of cuprizone exposure (**Figure 6G)**, the cells in L4 were primarily small and weakly fluorescent (**Figure 6F**), suggesting that they survived cuprizone treatment. Conversely, L6 contained bright oligodendrocytes with substantially larger somata (**Figure 6F and 7M**), indicating that they were recently generated. These results highlight how applying the ViT classifier as an added step in the analysis pipeline can distinguish between regions with high cell recovery and regions with high cell survival during injury. Detection of newly formed oligodendrocytes also provided a measure of the rate of change in oligodendrocyte density at any given timepoint. For instance, the high density of large oligodendrocytes in L6-SSp after cuprizone injury (**Figure 7M**) predicted the enhanced early recovery of L6-SSp three weeks post-injury (**Figure 6E**). Applying this predictive principle to datasets collected at three weeks of recovery suggests that L5-SSp, which exhibited the highest rate of oligodendrogenesis at this timepoint (**Figure 7N**), will recover the most in subsequent weeks. Together, these studies reveal the utility of ViTs, in combination with the overall clearing-imaging-analysis pipeline developed here, to identify sites of oligodendrogenesis in development, aging, and injury-repair contexts.

## DISCUSSION

Oligodendrocytes are solely responsible for generating myelin in the brain, which is critical for rapid sensation, information processing, and motor output. In addition to coordinating the timing of electrical impulses, which enables Hebbian learning through coincidence detection^78^, myelin provides metabolic support for underlying axons^79^ and regulates excitability by K^+^ clearance^80^. Although the density of myelin has been shown to vary between neuron types, brain regions, ages, and life experience, there has been no comprehensive brain-wide analysis of oligodendrocyte distribution across the lifespan to quantify these differences among functionally distinct brain regions. Here, we present a series of highly accessible and adaptable methodologies to determine the precise position of each oligodendrocyte in the murine brain. This approach, involving transgenic labeling, brain clearing, lightsheet imaging, image registration, and AI-based cell segmentation, allowed us to identify the position of over 300 million oligodendrocytes across 30 brains, providing information about regional control of oligodendrogenesis, variability in cellular patterning between individuals of different age, sex, and strain, and brain-wide differences in oligodendrocyte loss and regeneration in the context of injury. We demonstrated that oligodendrocyte distribution can be used to accurately estimate myelin coverage, overcoming technical challenges associated with quantifying the distribution and abundance of myelin. We extended these studies by developing a method to selectively identify newly formed oligodendrocytes within these large populations, showing how additional models can be applied to extract information from these terabyte-sized whole brain datasets. These studies provide new insight into the dynamic changes in oligodendrocyte patterning that occur within brain regions across the lifespan, and a series of computational tools that can be employed to study the regulation of developmental oligodendrogenesis, adaptive myelination with learning, and oligodendrocyte production in diverse physiological and pathological contexts.

The use of *Mobp-EGFP* mice enabled visualization of myelin sheaths due to the presence of EGFP in the myelinic channels (inner and outer tongue) and paranodal loops that extend along the length of each myelin sheath^51^. Oligodendrocytes extend cytoplasmic processes that branch to form ∼40-60 individual myelin sheaths^3,4^ within a local area surrounding the soma. In accordance, our results revealed that the area occupied by these processes was directly proportional to the number of oligodendrocyte somata; thus, oligodendrocyte soma analysis provides an accurate estimate of the amount of axonal territory covered by myelin within a given region. However, because EGFP is excluded from myelin itself, and the myelinic channel width is not proportional to myelin thickness^81^, this analysis does not provide information about myelin sheath thickness, a key feature of myelin that varies substantially between different axons and has been shown to be modified by behavioral experience in adults^82^. Implementation of other imaging modalities, such as reflected light (e.g. ScORe^30^) or third harmonic imaging^28^ that are sensitive to myelin thickness or using other genetic strategies to fluorescently label myelin itself, could be implemented to study this additional structural feature of oligodendrocytes in diverse contexts. Future models could also leverage additional biological/optical tools, such as a multi-fluorophore labeling approach (e.g. brainbow^83^), to resolve the complexity of densely myelinated tracts in white matter that can presently be disambiguated only through electron microscopy.

With speed, scalability, and image quality as guides, we chose to use lightsheet imaging of brains from *Mobp-EGFP* mice to map the distribution of oligodendrocytes. Adapting previously reported clearing methods, such as CUBIC-L^42^, that offered exceptional delipidation while retaining EGFP fluorescence, we resolved oligodendrocytes and individual myelin sheaths throughout the intact brain with high fidelity. The use of endogenous EGFP removed the need for immunostaining, which greatly extends the time needed for labeling, adds cost, and is prone to variable detection due to gradients in antibody penetration in thick tissues^84^. Immunolabeling at scale is also limited when assessing highly abundant antigens typically used for oligodendrocyte labeling, such as myelin proteins, due to antibody depletion^84^. Alternatively, serial block face two photon imaging has been used to define the morphology and distribution of fluorescent cells in the brain^34,52^; however, this approach requires specialized instrumentation and consistent serial sectioning, which is difficult to achieve and destructive. The gentle clearing pipeline we developed did not destroy endogenous fluorophores, and brains that were processed and imaged could later be re-imaged at higher resolution, or sectioned and analyzed with immunohistochemistry. While the average processing time (∼14 days) of our protocol was longer than other rapid clearing approaches^85^, fluorescent images from these brains had consistently high signal-to-noise required for subcellular resolution imaging, and batch processing was employed to clear groups of samples, reducing the overall time required for analysis. Future improvements could focus on improving clearing of tissue from older mice (> P640), which exhibited lower optical transparency in deep regions despite additional delipidation time and were more susceptible to other confounding sources of fluorescence, such as protein aggregates and blood vessels that required additional processing steps to avoid registration errors and cell detection artifacts.

To identify the location of the approximately 10 million oligodendrocytes in each mouse brain, we developed novel AI-based methods for cell segmentation that were accurate and consistent, allowing analysis of cohorts of animals of different age, sex, strain, and life experience. The deep neural networks we employed represent significant advances in the analysis of fluorescent BigData volumes. In particular, we adapted a 3D Mask R-CNN^48^ to work with volumetric fluorescent datasets, which required extensive hyper-parameter tuning and additional normalization layers, as most applications of Mask R-CNN are in 2D natural scene processing or detection of sparse objects in medical images^49^. Adapting the Mask R-CNN to our highly dense datasets required additional bounding-box sorting and splitting to accurately separate clustered cells. Moreover, as terabyte-sized datasets must be analyzed in patches, due to computational memory constraints, the bounding box segmentations from the Mask R-CNN allowed us to implement a sliding-window approach that could merge multiple detections at the edges of FOVs to create a seamless, accurate analysis of patched large datasets. We demonstrated that this sliding-window Mask R-CNN approach outperformed other popular CNN architectures, such as the UNet^46^, which for our application needed additional post-processing algorithms for cluster separation that can be difficult to optimize across datasets. Moreover, while alternative whole-brain cell detection algorithms often rely on sparsity or the specific compartmentalization of fluorophores in the cell soma to identify cells^86^, this is not possible in our datasets. As EGFP is expressed throughout oligodendrocyte processes and somata, a key feature of the Mask R-CNN was its ability to identify individual cells despite being surrounded by non-somatic EGFP fluorescence, such as in white matter. Finally, a major advantage of our deep learning models is their ability to generalize and improve over time. These flexible, fine-tunable models can be adapted to new conditions, recognize additional features, and overcome artifacts that manifest in particular experimental contexts.

After extracting individual oligodendrocytes from whole-brain datasets, registering their position to a common coordinate framework (CCF) is critical to enable comparisons of cell densities between different brain regions and distinct individuals. We selected readily available open-source platforms (BigStitcher, BrainReg) to help democratize our pre-processing pipeline, performing stitching, fusion, and CCF registration to generate detailed density maps of the murine brain. These comprehensive maps revealed that the oligodendrocyte content between distinct brain regions spans more than four orders of magnitude. This remarkable regional variation is particularly striking considering that OPCs, from which oligodendrocytes are generated, exhibit highly regular patterning and a similar density in the brain throughout adult life^5,59^. If OPCs have a similar inherent capacity to undergo differentiation, these results suggest that the diverse oligodendrocyte (and myelin) patterns are primarily established by factors in the local niche, such as the availability of myelin-compatible axons^20^, activity-dependent molecular signals^87^, or the proximal release of factors from neurons and other glial cells^88^. As these density differences were highly consistent between hemispheres, individuals, and sex, it suggests that oligodendrocyte patterning may endow specific properties crucial to the unique functions of these microcircuits. However, this oligodendrocyte patterning was not static, as oligodendrocytes continued to be generated in adulthood, with the density increasing more than 2-fold in many regions such as the motor and visual cortices. The rate of oligodendrocyte addition was proportionally similar between regions of the neocortex with age. Regions with few oligodendrocytes, like perirhinal cortex, remained sparsely myelinated, while the ratio of cell density between primary and supplemental areas was strikingly preserved throughout life in motor and somatosensory cortices, thereby maintaining the relative differences in oligodendrocyte density established early in life. The impact of this progressive, prolonged developmental myelination program on the functional properties of circuits is not yet known. However, as the rate of oligodendrogenesis can be enhanced acutely by learning^16^, sensory exposure^15^, or pathological conditions such as seizures^89^, and preventing oligodendrogenesis can impair motor learning^90^ and limit neuronal plasticity during visual development^91^, these profound developmental changes in myelin content are likely critical for shaping circuit function and behavior. Although the number of oligodendrocytes continued to increase in adulthood, the overall rate of oligodendrocyte addition slowed across all brain regions with age, reaching a near plateau after P620. Nevertheless, regional deviations from this global trend were observed, such as in layer 6 of the prefrontal cortex, where oligodendrocyte addition did not slow with age. Another deviation was in the cerebellar cortex, which plateaued much earlier in life, as few oligodendrocytes were added after P60, further highlighting the extreme regional control of oligodendrocyte patterning.

Oligodendrocytes within distinct brain regions also exhibited differences in their susceptibility to injury and regeneration that were highly consistent between individuals, with oligodendrocytes in the retrosplenial cortex and in layer 4 of sensory regions particularly resistant to cuprizone. Although local differences in oligodendrocytes between cortical regions have not been explored, single nucleus RNA sequencing studies indicate that the transcriptional state of oligodendrocytes varies substantially between gray and white matter^92^, which may result in differential sensitivity to this toxin and other insults. The recovery from demyelination was also variable, with deeper layers of motor and somatosensory cortex recovering exceptionally fast, despite also experiencing the most severe loss during injury. Recovery in regions closer to the subventricular zone may be facilitated by OPC production from *Gli1+* neural stem cells, which is enhanced by demyelination^93^. Expansion of our brain-wide analysis approach to simultaneously track the distribution of these progenitors and oligodendrocytes would help reveal the impact of this recruitment on regional oligodendrocyte regeneration. While it is possible that the destruction of oligodendrocytes merely turns back the developmental clock—freeing up territory along axons that are then remyelinated at the same relative rate as seen during earlier development (after accounting for age-dependent decline in OPC differentiation^94^)—our data suggests that regeneration is also strongly influenced by local features not present during development. For instance, after cuprizone injury, cortical and subcortical regions deviated from their earlier developmental trajectories. Thus, the dynamics of remyelination are likely influenced by both previously established cues (e.g. preserved axon characteristics) with varying features (e.g. neural activity patterns).

A significant advantage of our analysis pipeline is the ability to define the location of newly formed oligodendrocytes using their characteristic morphological features, providing information about rates of oligodendrogenesis from samples collected at a single timepoint. Applying this analysis, we found that oligodendrogenesis decreased globally across the lifespan but was still detected in animals over two years of age, indicating that the near-plateau of cell density with aging is in part due to a decline in oligodendrogenesis rather than an increase in turnover or death of oligodendrocytes. Moreover, by defining the distribution of newly formed oligodendrocytes, we were able to distinguish between brain regions that displayed high levels of recovery during injury, and regions that simply had a high density of oligodendrocytes that survived injury. Such analysis could be extended to determine the precise age of these cells and their probability for death, by expanding the model to use additional classifiers. However, classifiers dependent on fluorescence intensity and cell morphology are influenced by fluctuations in illumination and clearing homogeneity, necessitating high standardization between samples. Future ViT models for cell classification should aim to encode extraneous variables (laser power, exposure time, refractive index homogeneity) into model training, or design pre-processing steps to rigorously normalize across all datasets. At present, it is difficult to distinguish newly formed oligodendrocytes from clusters of mature oligodendrocytes in white matter or deeper brain regions where myelin density is high. Application of this cell mapping approach using additional transgenic reporter mice that specifically label newly formed oligodendrocytes^95^ would help reveal rates of oligodendrogenesis in these dense areas.

The results and tools compiled in this study serve as a new resource for comprehensively understanding the regional variability and dynamics of oligodendrocyte patterning in the murine brain. These datasets can be accessed using cloud-based tools, such as Neuroglancer, for individualized annotation and analysis, and both raw data and deep learning models are available to customize analysis through further training. These datasets are compatible with maps of brain vasculature^32^ and neurons^33^ referenced to the same atlas, facilitating exploration of the relationship between oligodendrocytes and blood vessels, which serve as a scaffold for migrating OPCs during development and during regeneration^96–98^, and specific neuron subtypes, such as interneurons, that are highly myelinated in the cerebral cortex^99^. By combining lightsheet, high resolution mapping of myelin sheath distribution with AAV-based sparse labeling of axons from distinct neuron types^34^, it will be possible to define the myelination pattern of distinct neuron subtypes brain-wide, generating a cell based “myelinome” to supplement emerging information about the connectome. This spatial information about neuron specific myelin patterning will aid in developing accurate models of brain circuits and enable assessment of changes to these patterns in health and disease to better understand the functional consequences of myelin plasticity and differential myelin repair.

## Supporting information

Supplemental Information (procedures)

## Acknowledgments

Funding was provided by the National Institute of Health, the Chan Zuckerberg Initiative, and the Dr. Miriam and Sheldon G. Adelson Medical Research Foundation. Y.K.X. is supported by a Johns Hopkins Kavli Neuroscience Discovery Institute Fellowship. We also thank A. Smirnov and M. Pucak for supporting the daily operation and maintenance of microscopes essential to the implementation of this study.

## Author contributions

Conceptualization, Y.K.T.X., J.S., D.E.B.; Methodology, Y.K.T.X., A.B., E.M.; Data curation and formal analysis, Y.K.T.X.; Funding Acquisition, D.E.B.; Investigation, Y.K.T.X., A.B., E.M., A.K., J.E.B.; Software, Y.K.T.X., E.M., S.Z.; Supervision, J.S., D.E.B.; Visualization, Y.K.T.X., E.M., A.K.; Writing – original draft, Y.K.T.X., D.E.B.; All authors contributed to review and editing the manuscript.

## METHODS

### Animal care and use

All animal experiments were performed in strict accordance with protocols approved by the Animal Care and Use Committee at Johns Hopkins University. Female and male adult mice were used for experiments and randomly assigned to experimental groups. All mice were healthy and did not display any overt behavioral phenotypes. Generation and genotyping of BAC transgenic lines from *Mobp-EGFP* (GENSAT^37^) have been previously described^15^. Mice were maintained in a climate-controlled room on a 12 hr light/dark cycle, housed in groups no larger than 5, and food and water were provided ad libitum, except during cuprizone-administration. All mice were maintained on a pure C57BL/6 (B6) background unless otherwise indicated as first generation mixed B6/FVB animals.

### Perfusion and tissue dissection

All mice were deeply anesthetized with sodium pentobarbital (100mg/kg w/w) prior to transcardial perfusion with 30mL PBS (pH 7.4) followed by 35 mL ice-cold 4% paraformaldehyde (PFA in 0.1 M phosphate buffer, pH 7.4). PBS and PFA volumes were doubled for older mice (> P620) to ensure blood clearance and sufficient fixation. The pH of all solutions is critical to retain optimal endogenous EGFP fluorescence. Brains were postfixed overnight at 4°C in 4% PFA and stored in PBS at 4°C before tissue clearing. Mice were perfused at P60 or as indicated.

### Cuprizone treatment

At 8 weeks of age, male and female *Mobp-EGFP* C57BL/6 mice were fed a diet of milled, irradiated 18% protein rodent diet (Teklad Global) supplemented with 0.2% w/w bis(cyclohexanone) oxaldihydrazone (cuprizone, Sigma-Aldrich) in custom gravity-fed food dispensers for six weeks. Treated mice were returned to regular pellet diet during a three-week recovery period.

### Tissue clearing pipeline

Whole brains were removed from PBS and transferred directly into fresh SHIELD OFF solution at 4°C (50% v/v SHIELD buffer, 25% v/v SHIELD Epoxy, in dH_2_O). All SHIELD reagents were obtained from Lifecanvas, and all steps were done with shaking on a nutator and covered from light. After 3 days of incubation with SHIELD OFF, the tissue was then transferred to SHIELD ON for overnight incubation at 37°C. After completion of the SHIELD protocol, whole brains were briefly passed through PBS before transferring to 50mL of CUBIC-L (10% v/v Triton X-100; 10% v/v *n*-butyldiethanolamine; 80% v/v dH_2_O) for delipidation. Samples were incubated at 37°C for a variable amount of time based on the age of the tissue (see Supplemental Information) due to increased lipid content in older tissue. CUBIC-L was refreshed every 3-4 days for the duration of delipidation. Notably, SHIELD protection reduced brain expansion but also altered the opacity of the brain post-CUBIC. With CUBIC-L alone, brains appeared translucent, but with SHIELD, they remained mostly opaque. This is expected and did not alter the clarity of imaging. After CUBIC-L, brains were washed with PBS for 1 hr followed by an overnight PBS wash at 37°C. Sample size, color, and opacity should be similar to unprocessed tissue before moving brains to the final clearing solution. For our clearing/imaging solution, we modified RIMS (∼60% w/v Nycodenz, 0.5% w/v Meglumine, 0.25% w/v Diatrizoic acid, and 0.01% sodium azide in 20mM PB buffer) by adding 40% w/v Urea and removing Tween20, as Tween caused tissue expansion and introduced bubbles into the cleared tissue that obstructed imaging. A high saturation of Urea was also critical in achieving optimal imaging at depth. Moreover, we adjusted the RI to ∼1.496 using a refractometer (Mettler Toledo Refracto 30GS). Increasing the RI further by adding more Nycodenz did not noticeably improve imaging quality but did increase the turbulence of the imaging solution as it neared saturation, which created random blur artifacts (**Figure S1F**). Finally, we found that altering the pH or solute content also impacted the degree of tissue expansion (higher pH causes more expansion), so we optimized RIMS for pH ∼7.4 to reduce expansion but also prevent EGFP degradation in an acidic pH. For detailed instructions on uRIMS formulation, see Supplemental Information.

Brains were transparent after 1 day in uRIMS but optimal quality required at least 2 days, especially for older tissue. We do not recommend storing brains in uRIMS indefinitely (+2 weeks) as the aqueous environment can begin to cause gradual tissue disintegration. Rather, we removed the brains entirely from uRIMS and stored our samples dry in air-tight 5 or 10mL tubes in the dark at room temperature until required for imaging, at which point we moved the sample back into uRIMS overnight at 37°C to re-equilibrate. Spinal cord samples were cleared in a similar fashion with slight modifications to incubation time at each step: 2 days SHIELD OFF at 4°C, 1 day SHIELD ON at 37°C, 3 days CUBIC-L at 37°C, 1 day PBS, 2 days uRIMS at 37°C.

### Lightsheet microscopy

All samples were imaged with a Zeiss Lightsheet 7 microscope. Rapid whole-brain scanning was performed with a 5× 0.16/NA air objective (ZEISS) and two 5× illumination objectives set orthogonal to the detection path. Whole brains were suspended vertically on custom 3D-printed holders using UV-activated optical glue (Ergo 8500), such that the dorsal surface faced the detection objective (**Figure 1E**). Sample holders were printed with black PLA (Ryno M-698-9PRZ) using a Prusa i3 MK3S+ 3D printer.

Acquisition tile FOV was a horizontal plane of the tissue with XY size of 1960 x 1960 pixels at a resolution of 1.15 µm/px with 10% overlap between neighboring tiles. Axial resolution was set at 5 µm/px and moved dorsal-ventral through the tissue. Overall, an entire brain could be imaged with a 6 x 7 (height x width) set of tiles, creating a volume of 10 mm x 12 mm x 9 mm (XYZ), and roughly 1 TB of size per imaging channel. As the Zeiss chamber has a maximum movement distance of 10 mm in XZ, it is exceptionally important to reduce tissue expansion during clearing.

To improve image quality due to light scattering, we used dual illumination imaging to ensure that each hemisphere of the brain was captured at least once while being illuminated with the objective most proximal to itself. Moreover, we found that high laser power (∼80%) with low exposure time (∼30 msec) provided optimal imaging resolution. We imaged *Mobp-EGFP* brains with simultaneous 488 nm and 638 nm lasers to excite EGFP and autofluorescence, respectively. For reduced bleed-through between channels, we used pre-set filter cubes to split the collected light (BP 505-545 + LP 660 emission filters). Acquisition time on average was about 4–6 hours per brain. Finally, we found that the addition of a layer of ∼3 mL sunflower oil (Sigma) pipetted onto the exposed surface of the imaging chamber greatly reduced evaporation and evaporation-induced turbulence artifacts, allowing for extended imaging sessions. Brains could be passed through the sunflower oil into uRIMS without creating artifacts, but removal of brains through the oil after imaging left residual oil residues. Thus, to re-image samples, we washed brains in PBS first to remove trapped oil before returning to uRIMS to re-equilibrate.

For high resolution imaging, we used either 20× Zeiss PN 20×/1.0 Corr nd=1.45 or PN 20×/1.0 Corr nd=1.53 immersion objectives, depending on the desired imaging RI, in addition to two 10X illumination objectives. The acquisition XY size was reduced (1960 x 1400 px) to remove optical aberrations at the edges of each tile. Each FOV was acquired at a resolution of 0.2 um/px in XY and 1 um/px in Z with 10% overlap between neighboring tiles. Due to the optical limitations of the high-resolution objectives, the maximum whole tissue that could be fit into an imaging FOV was 1 full hemisphere at a time. For a whole brain, we estimated the data size to be ∼30 TB and would require 2-3 consecutive days of imaging. All filter sets and sample mounting techniques were the same for high- and low-resolution data.

### Data pre-processing and stitching

Dual-channel, dual-illumination raw CZI files with 10% overlap (∼2TB, ZEISS) were first converted to hdf5 file format using BigStitcher (FIJI) with deflate compression (∼1TB). Illumination selection was done in BigStitcher to divide the data in half sagittaly, where the illumination side for each tile was assigned to the closest respective illumination objective. We then used BigStitcher to perform interest-point based affine image stitching to align adjacent tile overlaps. Interest points were first detected in the oligodendrocyte (*Mobp-EGFP*) channel as cell bodies. We then used either “Fast descriptor-based (rotation invariant)” or “Precise descriptor-based (translation invariant)” algorithms to perform interest-point based stitching. Optionally, if chromatic aberrations persisted between imaging channels, we employed ICP refinement at 8x resolution which sometimes could improve the alignment between imaging channels. Data was then fused and saved in the N5 file format which is accessible from Python through lazy-loading and is compatible with Neuroglancer for data sharing. All steps were performed on an Ubuntu computer with an i9 20-core/40-thread processor with a 4070 GPU and 256 GB RAM. Additionally, prior to analysis, we also manually curated the dataset to exclude regions that were torn or obstructed by bubbles as well as those that were hazy due to poor clearing (**Table S1**).

### Brain registration to CCF

All registration steps were performed on downsampled (20 µm/voxel) isotropic volumes of the acquired autofluorescence channel (638 nm excitation), unless otherwise indicated. Prior to registration, each volume was first enhanced with adaptive histogram equalization, followed by stripe filtering (FIJI^100^, Xlib plugin, settings: wavelet-FFT, both directions, wavelet=Sym5, decomposition=0:5, damping=2) to remove tiling illumination artifacts, and N4 bias correction^101^ to reduce uneven illumination (**Figure S3A**). Registration was then performed using the BrainReg platform^55^ at 20 µm/px resolution (grid spacing=-10, bending energy=0.9). Notably, diffeomorphic registration with ANTS^57^ was visually more appealing, but resulted in excessive warping when examining myelin patterns. For instance, layer 2/3 oligodendrocytes were warped into layer 1 (**Figure S3B**).

One common complication for registration is the mismatch in autofluorescence that arises from different clearing techniques and imaging modalities. For instance, since the Allen CCF autofluorescence was collected using a serial two-photon microscope, it differs in significant ways from autofluorescence maps captured with iDISCO or CUBIC lightsheet imaging pipelines (**Figure S3C**). In our case, we noticed that the most substantial difference was in regions with high myelin content, such as white matter tracts, whereby the Allen CCF was considerably darker than our autofluorescence fiber tracts. Serendipitously, as we were imaging myelin in the *Mobp-EGFP* channel, we could identify white matter tracts easily by thresholding, and then divided our autofluorescence channel by the masked white matter regions to produce darker fiber tracts that more closely resembled the Allen CCF (**Figure 3**). This improved white matter registration substantially (**Figure S3D,E**) and allowed us to generate our own average template maps that removed the reliance on myelin division for future experiments that lacked *Mobp-EGFP* (**Figure S3H**). A similar division-based approach was used to subtract blood vessel artifacts, which would often appear in brains that were not optimally perfused. As blood autofluorescence would appear in both 488 and 638 nm channels, we could first divide our 638 nm autofluorescence channel by the 488 nm EGFP channel to create a volume with amplified blood artifacts. We then divided our 488 nm EGFP volume with the amplified blood vessel volume to dampen vessel artifacts.

An additional complication was the appearance of autofluorescent “blobs” in aged tissue. These blobs were only excitable by the 638 nm laser, which prevented them from bleeding-through and causing false positives in the EGFP channel, but they still served as significant sources of variability for registration. Notably, these blobs never overlapped with oligodendrocyte somata, and appeared to align with neuronal cell somata (**Figure S3I**). Thus, they are likely metabolic by-products that accumulate with age and not simply lipofuscin. These neuronal blobs motivated the establishment of an average myelin template, which was the mean of all registered *Mobp-EGFP* brains at P60. Older tissue (> P620), with significant neuronal blobs, were thus registered to the average myelin template from P60 rather than the autofluorescence template, to avoid registration errors created by severe blob autofluorescence.

Finally, we also noted that due to our UV-glue based mounting method, some brains would bow-out near the cortex/cerebellum border, causing these two regions to touch. Unfortunately, registration algorithms have a difficult time creating a gap when there is none to begin with, and thus would create distortion artifacts along this edge. To resolve this issue, we segmented the cerebellum using an overfitted ANTS transform, and then registered the cerebellum separately from the rest of the brain (**Figure S3G)**.

### Data visualization for density maps and flatmounts

Two types of density maps were created for visualization. Local summation-based maps were created by expanding the centroid of each detected cell into a spherical neighborhood with a radius of 350 µm. All neighborhoods were then summed together to generate local density counts for most figures. For analysis of large newly formed oligodendrocytes, however, the number of cells was too low for local summation-based maps. Thus, we simply calculated the density of oligodendrocytes per CCF region and re-plotted the CCF with the associated density values, creating region-based density maps.

For flatmount representations, we used the CCF streamlines package and associated streamlines calculated by the Allen institute^52^. Since oligodendrocyte patterns are unique between cortical layers, we could not simply create a maximum projection across all layers. Instead, we masked out each cortical layer individually prior to flatmount projection, thus creating layer-specific flatmounts.

### Mask R-CNN for instance segmentation of oligodendrocytes

To extract volumetric segmentations of oligodendrocyte somata in sparse and dense regions across the brain, we adapted and trained a 3D Mask R-CNN model^48,49^. Input volumes (128 x 128 x 16 voxels), containing *Mobp-EGFP* labelled oligodendrocytes, were cropped from high-resolution whole-brain datasets using a sliding window approach (**Figure 2B**). A total of 5,482 training and 685 validation volumes were cropped from across eight brains. Input volumes were then passed to a feature extraction network, built on a ResNet-50^102^ backbone, which used convolutions to extract features from a variety of scales. A region-proposal network (RPN) then convolved the extracted feature maps to propose regions-of-interests (ROIs) which were pooled together using ROIAlign (**Figure 2A**). Pooled ROIs and proposed anchors were then sent to both the fully connected classifier head, which classified each proposed anchor as either background or signal, and the segmentation mask head, which created a binary mask for the actual location of the identified object.

Post-processing was minimal, to reduce the number of hyper-parameters required for tuning the analysis pipeline. The primary source of error that required post-processing was the edge-effect, whereby detections made on the edges of a cropped FOV were likely to be partial or incorrect due to insufficient data for extrapolation. Thus, we implemented a size restriction, such that only objects > 30 voxels were considered as potential cells. As well, we used weighted-box clustering^50^ to accumulate detections across multiple FOVs that were analyzed with 10% overlap in XY and 20% overlap in Z, thus each “edge” detection would be centered in at least one FOV and altogether, each cell would be detected several times and these detections would be pooled together using WBC (**Figure 2B**). Finally, because bounding boxes denote a cube-shaped region of interest, they can overlap with one another unintentionally, due to protruding corners of the cube. Thus, we implemented “box splitting” to divide overlapped areas and assign them to a specific box, which allowed us to use the boxes to mask out individual cells in the binary segmentation mask (**Figure 2C**). Box splitting was implemented simply by finding the box centroids, and then assigning voxels bounded by all boxes to the nearest-neighbor box centroid.

### UNet myelin and cell segmentation

Two UNet models^46^ with 3D convolutional kernels were built in PyTorch 2.1.0^103^ for segmentation of myelin and oligodendrocyte somata, respectively (**Figure S2B**). Both CNNs contained four down-sampling convolutional blocks with 5 × 5 × 5 filters, batch normalization, and max pooling to extract local features. Four upsampling operations were then performed in reverse using trilinear upsampling and 1 × 1 × 1 convolutions to resize the image back to the same input size. A final 1 × 1 × 1 convolution reduced the output to a two-channel volume which was softmaxed with a threshold of 0.5 to two classes corresponding to background and cell soma (or myelin). Training was performed using a batch size of 8 for 120 epochs for myelin segmentation and 175 epochs for oligodendrocyte segmentation on an RTX 3070 GPU. Loss was calculated as cross entropy and optimized using an Adam optimizer with weight decay^104^ set at a learning rate of 10^–5^. Input volumes were 128 × 128 × 16 voxels and consisted of 3,295 training/367 validation images for myelin segmentation and 5,550 training/617 validation images for oligodendrocyte somata segmentation. All images were normalized by subtracting mean and dividing by standard deviation during training and inference.

### Vision Transformer cell classifier

We employed a custom Vision Transformer (ViT) model to classify oligodendrocytes as either newly formed or mature. The base ViT was pre-trained with 14 million images from ImageNet-21k^105^. Each volumetric input image (80 x 80 x 16 voxels) was centered around an oligodendrocyte of interest and represented as an 80 x 80 image with 16 input channels. We trained the model using binary cross-entropy with AdamW optimizer at a learning rate of 0.001 with scheduled rate reduction by 0.1 multiplier after plateau of 30 epochs. The training set consisted of 6,845 images with an additional 1,712 images for validation. Class balancing was implemented to ensure that all datasets contained 80% of class 0 (mature oligodendrocytes) and 20% of class 1 (newly formed oligodendrocytes). All images were normalized by subtracting mean and dividing by standard deviation. Models were trained for 200 epochs and the best model was selected based on validation loss and accuracy. All ViT models were implemented in Pytorch using the HuggingFace library^106^ and a Quadro RTX 5000 GPU.

### Adipo-Clear modified iDISCO clearing protocol

To compare to immunolabeling-based clearing methods, we applied the modified iDISCO method^40^ described as Adipo-Clear^107^ to *Mobp-EGFP* brains. All steps were performed while shaking on an orbital shaker or nutator. In brief, samples were delipidated in 30 min intervals at 37°C using an increasing methanol concentration at 20%, 40%, 60%, 80%, and 100% in B1n buffer (0.3M glycine, 0.1% (v/v) Tritonx-100, 0.01% (w/v) sodium azide). Samples were then incubated in dichloromethane (DCM) followed by a decreasing methanol gradient of 100%, 80%, 60%, 40%, and 20% in B1n buffer. Samples were washed in PTxWH (1x PBS, 0.1% (v/v) Triton x-100, 0.05% (v/v) Tween-20, 2µg/ml Heparin, 0.01% (w/v) sodium azide) before labeling with immunohistochemistry. Samples were incubated in primary antibody (Anti-GFP, Aves, RRID: AB_2307313) for 7-14 days followed by an incubation in secondary antibody for 7-14 days. Samples were washed in PTxWH then PBS before being immersed in an increasing methanol gradient of 25%, 50%, 75%, and 100% in H_2_O. Afterwards, samples were immersed in DCM for at least 1 hour at room temperature before being incubated in dibenzyl ether (DBE) for refractive index matching. Samples were incubated at least 1 day prior to imaging. Evaluation of antibody penetration was achieved by cutting the sample coronally and then imaging the outwards facing side of the sliced tissue.

### SHIELD-Delipidation-EasyIndex clearing protocol

To compare with commercially available clearing solutions, we applied the Passive Clearing Kit (LifeCanvas Technologies) to post-fixed *Mobp-EGFP* mouse brains. All steps were performed while shaking on an orbital shaker or nutator. SHIELD OFF solution was prepared fresh with DI water, SHIELD Buffer, and SHIELD Epoxy in a 1:1:2 ratio. Samples were incubated in SHIELD OFF for three days at 4°C, followed by SHIELD ON incubation at 37°C for 1 day. Samples were then delipidated with Delipidation Buffer at 37°C for 7-10 days, depending on the age of the mouse tissue. Delipidated brains were then rinsed with PBS before immersion in 50% EasyIndex in water overnight followed by 100% EasyIndex for refractive index matching. Samples were incubated for at least 2 days prior to imaging. In all comparisons, we observed that substituting CUBIC-L for Delipdation buffer improved quality of delipidation (**Figure S1D**) with some minor increase in tissue expansion (**Figure S1E**). We also noticed no obvious differences in image quality in using EasyIndex over uRIMS, however, notably, EasyIndex was less prone to crystallization with exposure to air, although EasyIndex encountered similar turbulence-based hazy artifacts (**Figure S1F**) likely due to the high saturation and RI of EasyIndex.

### Post-Clearing immunohistochemistry

Cleared tissue (uRIMS) was washed in 20mM PB buffer for 24 hours at room temperature before it was immersed in 30% sucrose (in PBS, pH 7.4) at 4° C for 24-48 hours, or until it was no longer floating. Tissue was then embedded in O.C.T (TissueTek 4583, Sakura) before it was cryosectioned into 50 µm slices (Leica CM1860 cryostat). Sections were collected in 20mM PB buffer with 0.1% NaN_3_ and kept at 4°C until use. For immunohistochemistry, sections were washed in 20mM PB buffer before permeabilization in 0.5% Triton x100 in 20mM PB buffer for 10 mins, followed by incubation with blocking solution (10% Normal Donkey Serum, 0.3% Triton X100 in 20mM PB buffer) for 1 hr. All steps were performed at room temperature with shaking. Samples were then incubated in primary antibody for 24 hrs at room temperature, washed 3 times with 20mM PB buffer, and incubated with secondary antibody for 24 hours at 4°C. After staining, the tissue was washed 4 times with 20mM PB buffer and sections were mounted onto slides with Aqua Polymount (18606, Polysciences) and allowed to dry for 24-48 hours at 4°C prior to imaging. The following primaries were used in this study: Anti-GFP (Aves, RRID: AB_2307313), Anti-Aspartoacylase (Genetex, RRID: AB_2036283), Anti-CD31/PECAM-1 (R&D Sytems, RRID: AB_2161028), Anti-NG2 (Bergles lab), Anti-Olig2 (Millipore Sigma, RRID:AB_570666). Images were acquired using a brightfield Keyence BZ-X710 at low magnification (5× objective), and with a Zeiss LSM 800 confocal at higher magnification (Zeiss 20× air and 40× oil objectives).

### Standard immunohistochemistry

PFA fixed tissue was embedded in O.C.T (TissueTek 4583, Sakura) before it was cryosectioned into 50 µm slices (Leica CM1860 cryostat). Floating tissue sections were washed in PBS before blocking (10% Normal Donkey Serum, 0.3% Triton X100) for 1 hr. Samples were then incubated with primary antibodies for 1 hr, followed by three more washes and incubation with secondary antibodies overnight. All steps were performed at room temperature with shaking. The tissue was then mounted onto slides with Aqua Polymount (18606, Polysciences) and imaged with a Zeiss LSM 880 confocal microscope (Zeiss 40× oil objective).

### Data and code availability

The datasets from the current study are available through Neuroglancer. All code and links to shared raw data can be found at https://github.com/yxu233/Xu_Bergles_brainwide_oligo_map

### Statistical analysis

All statistical analysis was performed using Python statsmodels and scipy libraries. *n* represents the number of animals used in each experiment, unless otherwise noted. Data are reported as mean ± s.e.m. or and *p* < 0.05 was considered statistically significant.

## SUPPLEMENTAL FIGURES

**Supplemental Figure 1.**
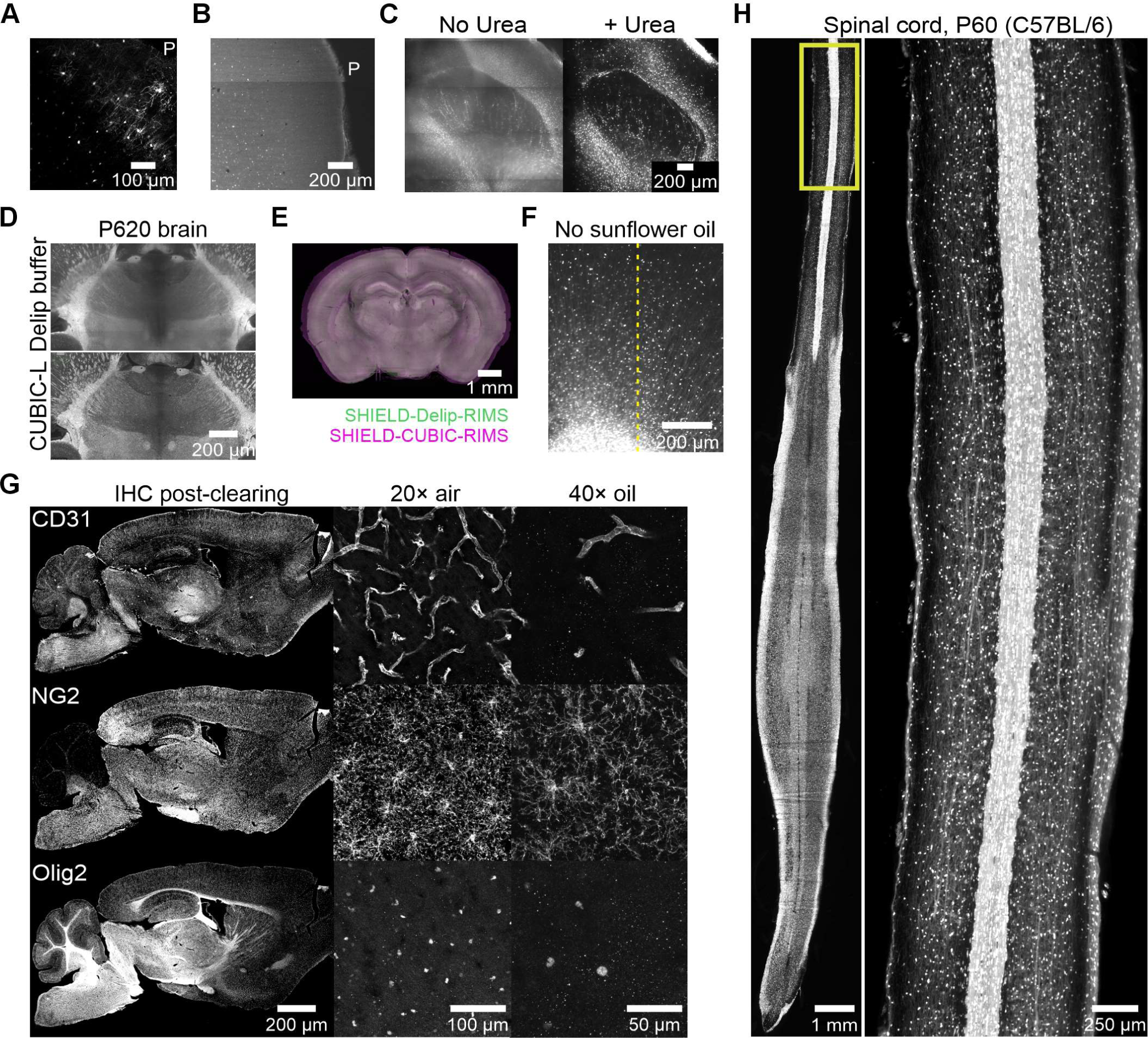
Optimization of clearing methods for endogenous fluorescence. A. Anti-GFP antibody penetration is poor in whole *Mobp-EGFP* brain cleared with Adipo-Clear. “P” denotes pial surface with loss of immunostaining in deeper layers. B. Organic solvents degrade endogenous EGFP fluorescence. “P” denotes pial surface with some preservation of endogenous EGFP in deeper layers. C. Clearing quality in deep striatal regions with and without 40% Urea in RIMS. D. Comparison of clearing quality with Delipidation Buffer (LifeCanvas) and CUBIC-L in old tissue. E. Tissue expansion is similar between SHIELD treated brains processed with Delipdation Buffer and CUBIC-L. F. Example of spontaneous dis-equilibration artifact when no sunflower oil is used during imaging. Left of dotted line is blurry and right of dotted line is clear. G. IHC with CD31, NG2, and Olig2 antibodies after tissue is cleared and then recovered by washing off uRIMS. 50 µm cryo-sectioned slices shown at different magnifications. H. Whole-mount 5× lightsheet imaging of a cleared spinal cord.

**Supplemental Figure 2.**
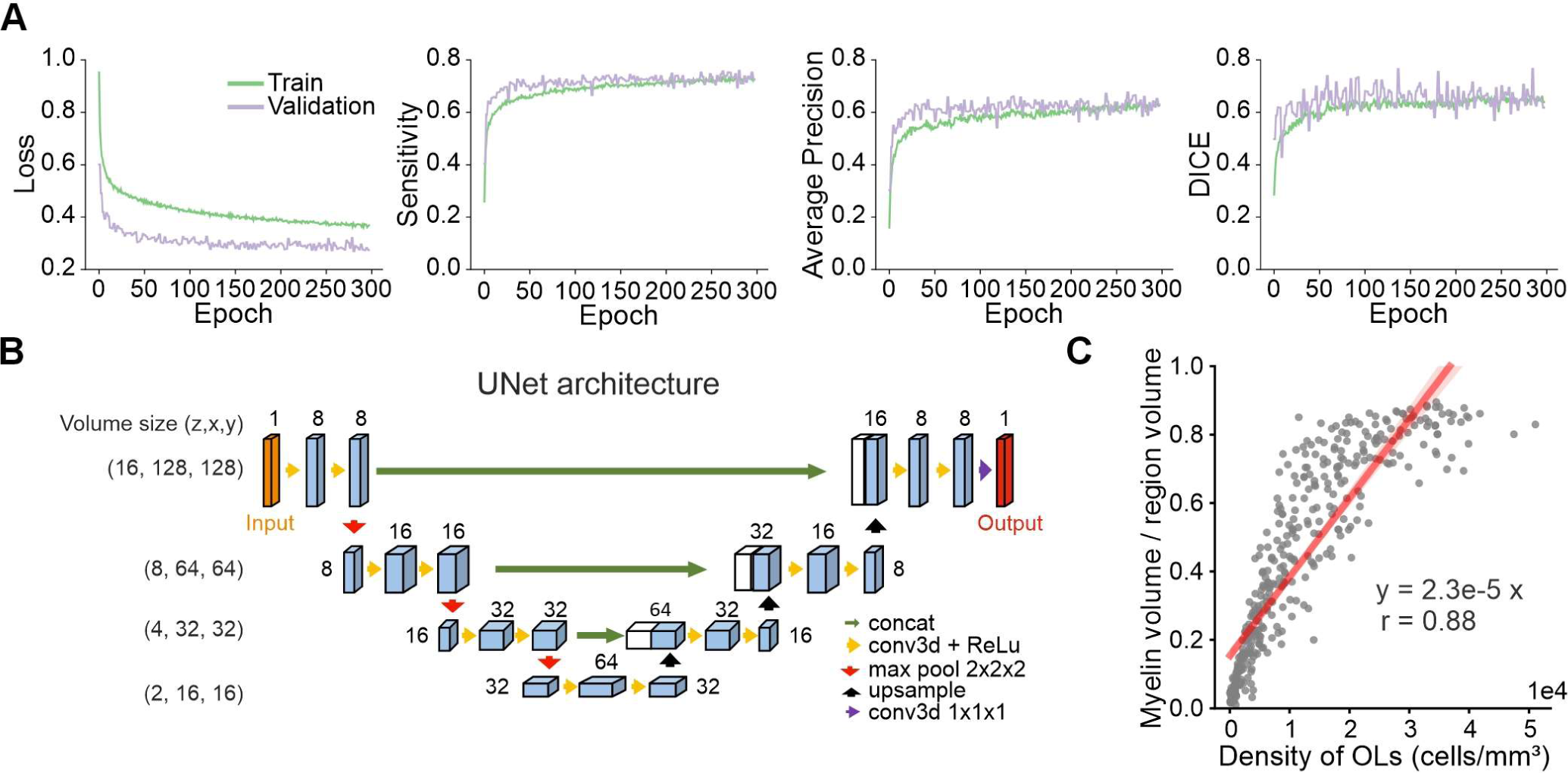
Performance of Mask R-CNN and UNet models. A. Training and validation curves for Mask R-CNN. B. UNet architecture used for myelin segmentation and oligodendrocyte cell body segmentation. C. Relationship between absolute myelin volume and oligodendrocyte density across CCF regions (r=0.88, *p=*2.9e-131, n=402 regions).

**Supplemental Figure 3.**
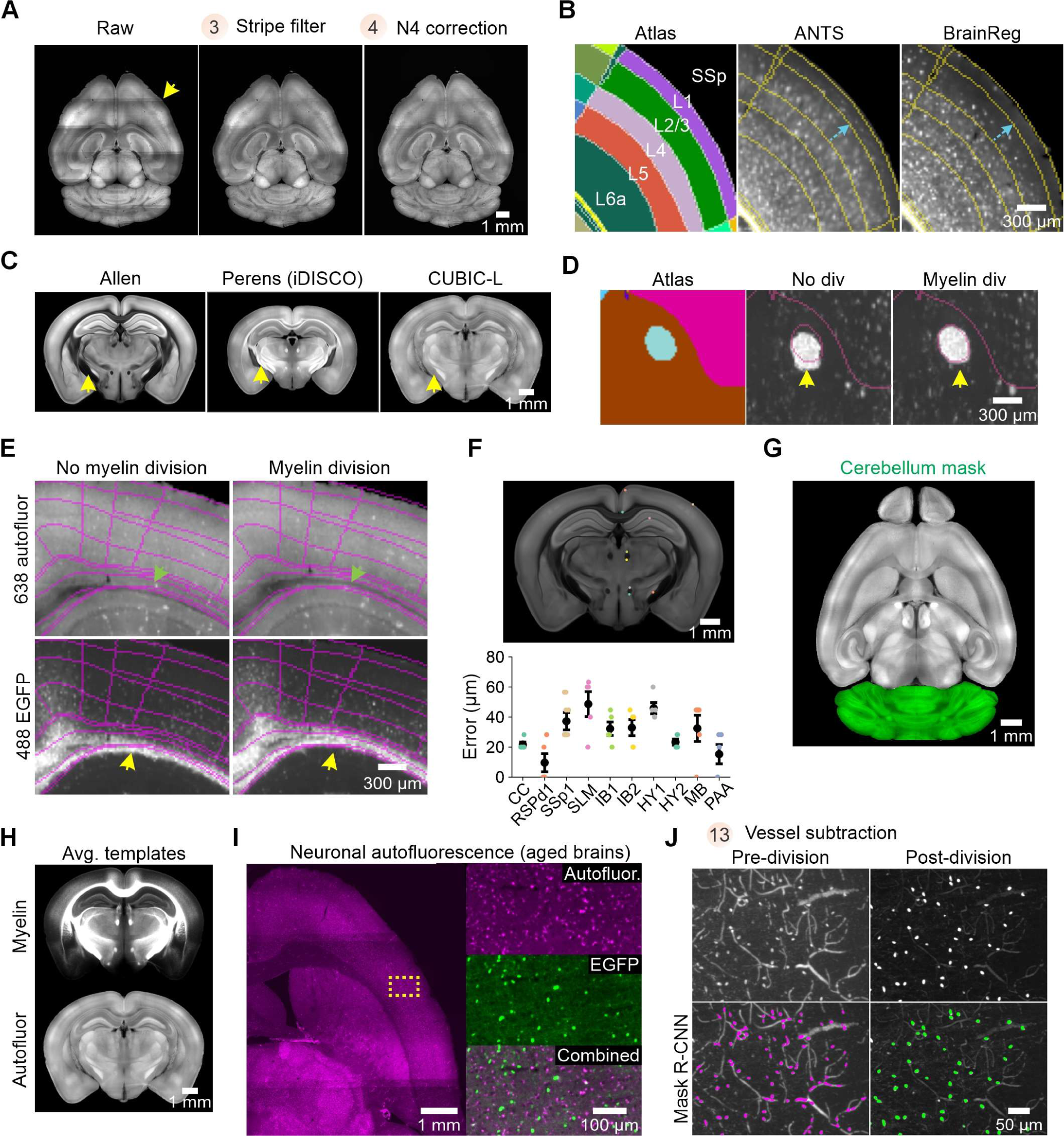
Additional pre-processing steps for CCF registration and analysis. A. Pre-processing steps for downsampled autofluorescence prior to registration. Stitch lines (yellow arrow) removed by stripe filter. B. Comparison of tissue warping with ANTS and BrainReg registration. Blue arrow depicts heavy warping of ANTS registration, where L4 oligodendrocytes are shifted into L2/3. This warping is not observed with constraints set in BrainReg. C. Single plane from Allen CCF autofluorescence, Perens atlas^108^, and our CUBIC-L autofluorescence, to depict differences in autofluorescence signal. Most striking contrast is in white matter (yellow arrow), where Allen atlas is exceptionally dark. D. Registration of fiber tract before and after myelin division to correct for autofluorescence mismatch. E. Registration of callosal fiber tracts before and after myelin division. F. Validation of registration accuracy using fiducial landmarks and corresponding measure of error (below). The atlas resolution is 20 µm/px, so an error of 1 pixel is equivalent to a 20 µm shift. G. Segmentation of cerebellum (green) from cortex facilitates accurate registration along tissue edge. H. Average autofluorescence and myelin templates created from our registered brains. I. Neuronal autofluorescence identified in aging tissue. No bleed through into EGFP channel (right inset). J. Mask R-CNN detections before and after vessel dampening by dividing raw EGFP volumes by blood vessel autofluorescence to mask out dual-channel bleed through.

**Supplemental Figure 4.**
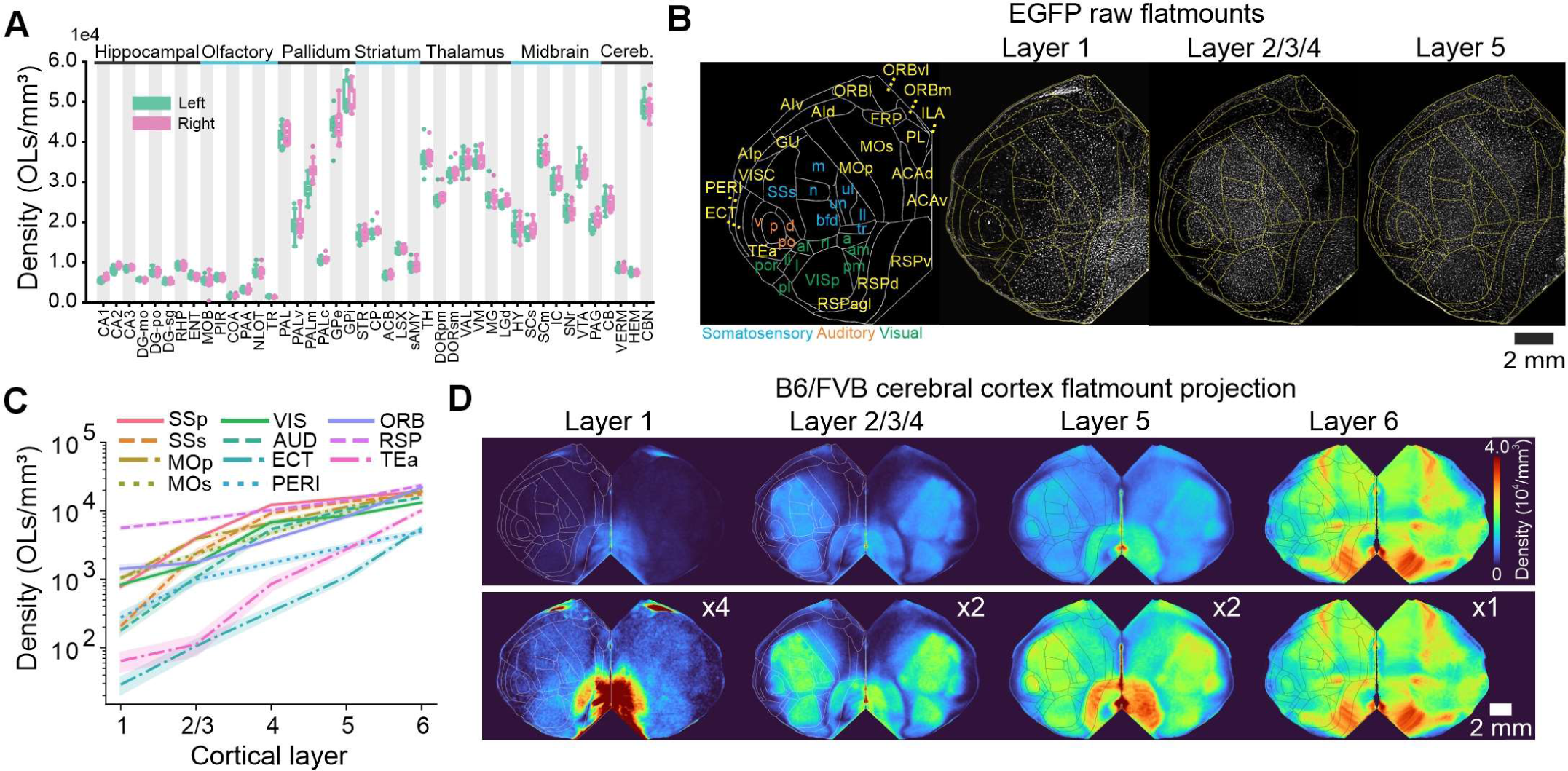
Oligodendrocyte density across the brain. A. Density of oligodendrocytes across select brain regions. B. Cortical flatmounts of raw *Mobp-EGFP* fluorescence from 1 mouse. C. Logarithmic-scale graph highlighting oligodendrocyte density across layers and regions of neocortex. D. B6/FVB cortical flatmounts of average cell density maps across cortical layers scaled (top) and unscaled (bottom).

**Supplemental Figure 5.**
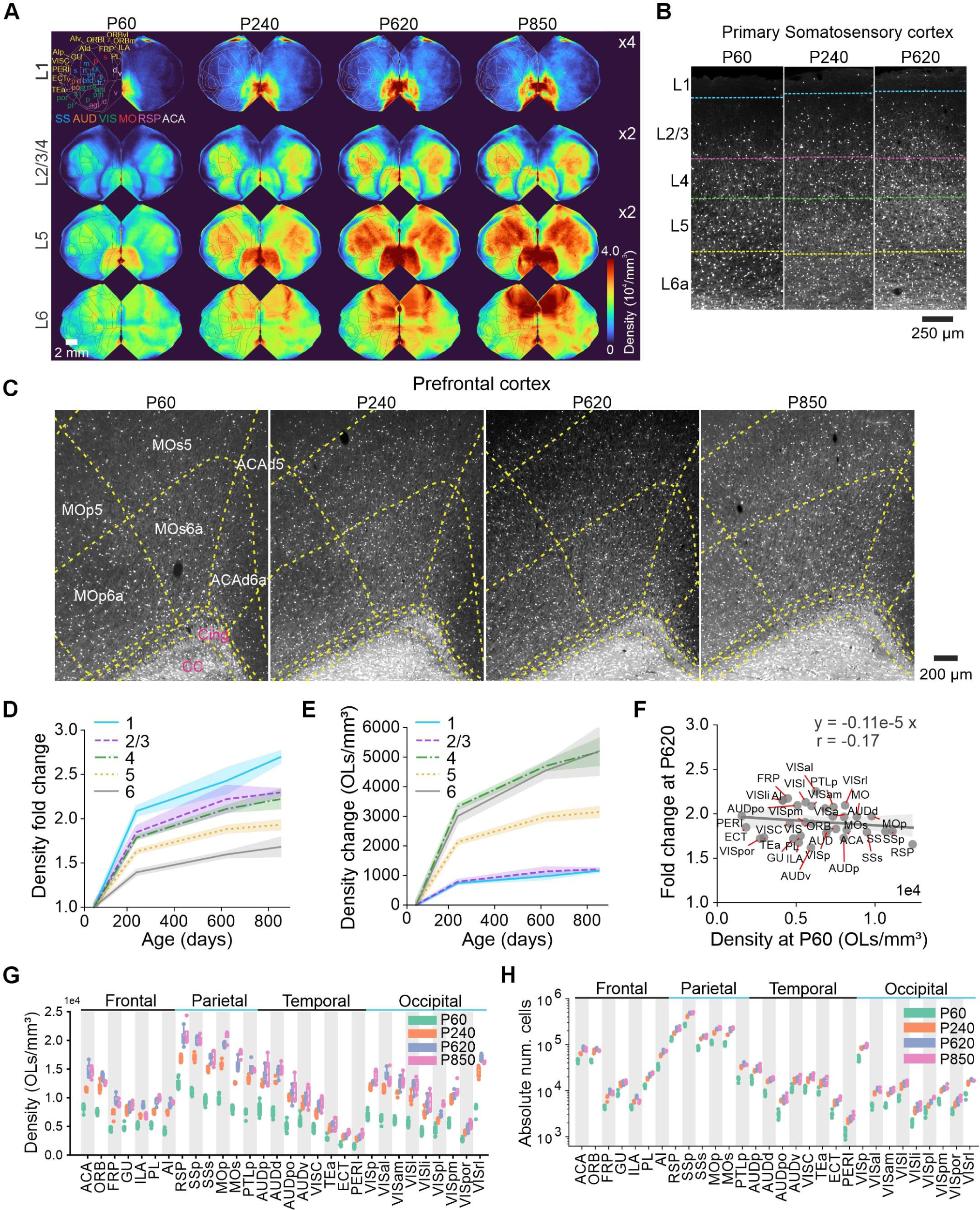
Layer-specific changes in oligodendrocyte density with age. A. Scaled cortical flatmount projections of average cell density maps to better visualize the distribution of oligodendrocytes in sparsely myelinated superficial layers. B. Example raw *Mobp-EGFP* data from primary somatosensory cortex across aging timepoints, C. and prefrontal cortex. D. Normalized trends of cortical layer-specific oligodendrocyte density pooled across all regions (n=8 for P60 and n=4 for all other timepoints). E. Absolute change in density of cortical layer-specific oligodendrocyte density pooled across all regions (n=8 for P60 and n=4 for all other timepoints). F. No correlation between oligodendrocyte density at P60 and normalized fold change at P620 (r=-0.17, *p=*0.35, n=34 regions). G. Oligodendrocyte density for all regions of neocortex at each aging timepoint. H. Raw absolute number of oligodendrocytes per region regions (n=8 for P60 and n=4 for all other timepoints).

**Supplemental Figure 6.**
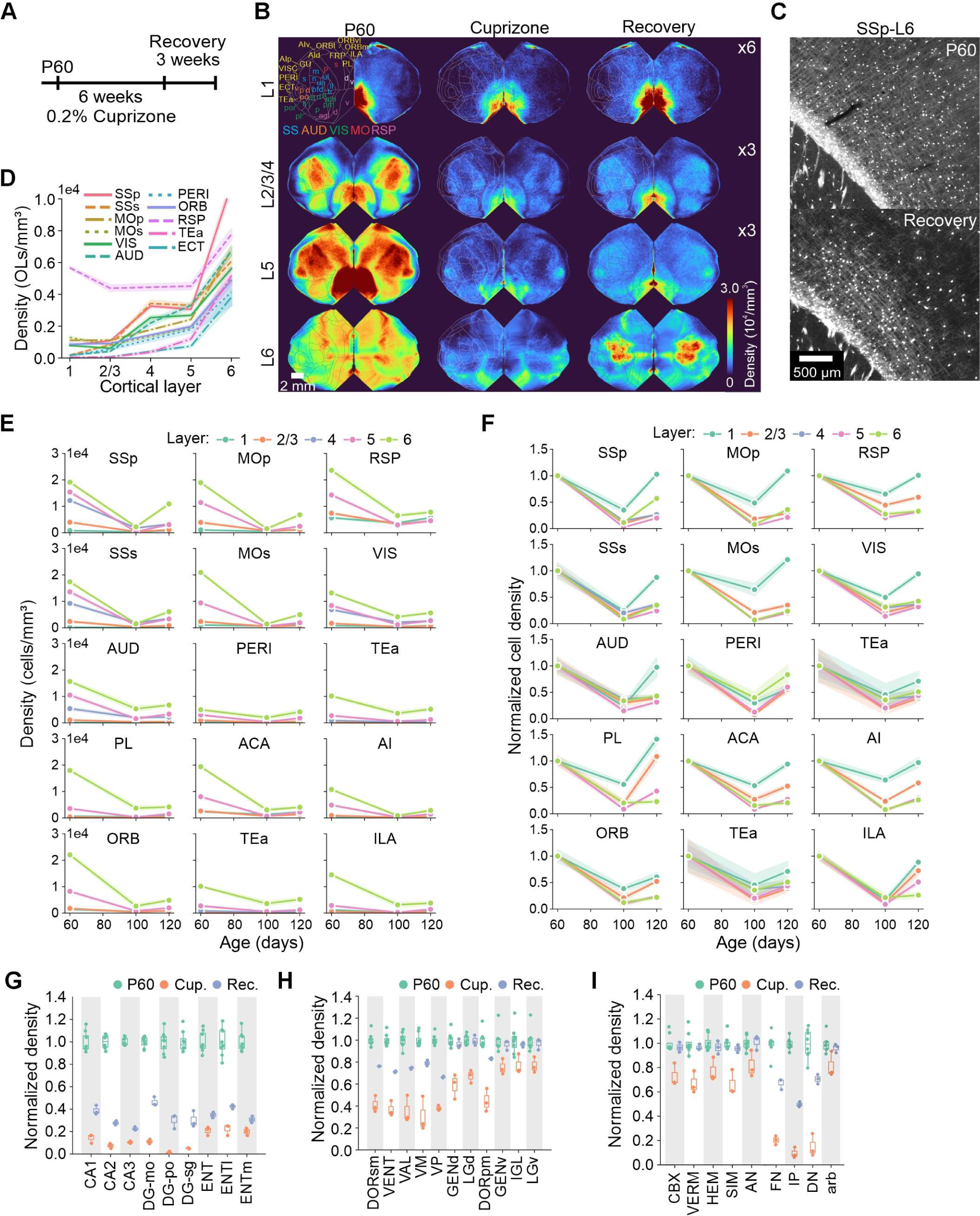
Layer-specific changes in oligodendrocyte density after cuprizone. A. Schematic of cuprizone injury-recovery experimental paradigm. B. Scaled cortical flatmount projections of average cell density maps to better visualize the distribution of oligodendrocytes during injury when cell density is low. C. Cyan inset from Figure 6C, showing laterally and vertically myelinated fiber tracts at baseline (P60) and during recovery. D. Density of oligodendrocytes across layers and regions during recovery. E – F. Absolute change (E) and normalized fold change (F) in oligodendrocyte density across cortical regions and layers from baseline to recovery. G – I. Normalized fold change from baseline to recovery for hippocampus (G), thalamus (H), and cerebellar cortex (I).

## SUPPLEMENTAL TABLES

**Supplemental Table 1:** Metadata for all brains analyzed and regions excluded.

| Mouse number | Experimental group | Exact age (P) | Weight (g) | Sex | Delipidation reagent | Myelin thresh | Blood div | Excluded CCF regions (regions – hemisphere) |
| --- | --- | --- | --- | --- | --- | --- | --- | --- |
| M127 | P60 | 62 | 20 | M | Delip Buffer | 500 | N |  |
| M229 | P60 | 58 | 20 | F | Delip Buffer | 450 | N |  |
| M126 | P60 | 62 | 19 | M | Delip Buffer | 500 | N | PL, ORB, ILA, FRP, ACA – (Left) |
| M223 | P60 | 59 | 25.2 | M | CUBIC-L | 400 | N |  |
| M299 | P60 | 58 | 25.5 | M | CUBIC-L | 500 | N |  |
| M254 | P60 | 58 | 16 | F | CUBIC-L | 350 | N |  |
| M256 | P60 | 58 | 17.5 | F | CUBIC-L | 350 | N |  |
| M260 | P60 | 56 | 18 | F | CUBIC-L | 500 | N |  |
| M279 | P240 | 242 | 34.5 | M | Delip Buffer | 500 | N | STR, TH, HY, IG, IB CB - (Both) |
| M286 | P240 | 243 | 29 | F | CUBIC-L | 400 | N |  |
| M281 | P240 | 242 | 33 | M | CUBIC-L | 400 | N |  |
| M285 | P240 | 243 | 25.5 | M | CUBIC-L | 400 | Y | CB – (Both) |
| M271 | P620 | 616 | 26 | M | CUBIC-L | 750 | N |  |
| M91 | P620 | 662 | N/A | M | Delip Buffer | 450 | Y | Torn areas (PL, ORB, ILA, FRP, ACA, CB – Both) |
| M97 | P620 | 624 | N/A | M | CUBIC-L | 400 | N | Right hemisphere |
| M334 | P620 | 620 | 36.6 | F | CUBIC-L | 500 | N | CB – (Left) |
| 1Otx7 | P850 | 856 | 34.1 | F | CUBIC-L | 500 | Y | Missing OB causing artifacts (PL, ORB, ILA, FRP, ACA, MB, IB, CB - Both) |
| Otx18 | P850 | 800 | 26.6 | M | CUBIC-L | 300 | Y | MB, IB, CB – (Both) |
| 5Otx5 | P850 | 858 | 31.4 | F | CUBIC-L | 300 | Y |  |
| Otx6 | P850 | 956 | N/A | F | CUBIC-L | 500 | Y |  |
| M147 | FVB | 58 | 18 | F | Delip buffer | 450 | N | RSP – (Left)<br>HY, MED, Vn – (Both) |
| M155 | FVB | 59 | 26 | M | Delip buffer | 400 | N | RSP – (Left)<br>HY, MED, Vn – (Both) |
| M152 | FVB | 59 | 22.4 | M | Delip buffer | 500 | N | RSP – (Left)<br>HY, MED, Vn – (Both) |
| M146 | FVB | 58 | 18.1 | F | Delip buffer | 500 | Y | HY, MED, Vn – (Both) |
| M265 | Cuprizone | 101 | 24 | M | CUBIC-L | N/A | N |  |
| M266 | Cuprizone | 101 | 22.5 | M | CUBIC-L | N/A | N |  |
| M267 | Cuprizone | 101 | 22 | M | CUBIC-L | N/A | Y |  |
| M312 | Cup-Recovery | 120 | 18.4 | F | CUBIC-L | N/A | N |  |
| M310 | Cup-Recovery | 120 | 20 | M | CUBIC-L | N/A | N |  |
| M313 | Cup-Recovery | 120 | 21 | F | CUBIC-L | N/A | N |  |
| Excluded from all brains | Hypothalamus – PVZ, PVR, MBO, RCH, VMH, ME<br>Hindbrain – All regions<br>Midbrain – MBsta<br>Cortical subplate – BMA, PA |  |  |  |  | Thalamus – Xi<br>Midbrain – SCO<br>Cerebellum – LING<br>Fiber tracts – cm, pyd, cbp |  |  |

## SUPPLEMENTAL VIDEOS

High-resolution videos can be found here: https://vimeo.com/1003497448?ts=0&share=copy

**Video 1. Whole brain volume of oligodendrocyte patterns**. Fluorescent *Mobp-EGFP* mouse brain imaged with ZEISS lightsheet microscope using a 5×/0.16 NA objective. Volume size 12 × 14 × 8 mm (XYZ).

**Video 2. Cross-section of myelin patterns from motor cortex to hypothalamus**. Cropped FOV from fluorescent *Mobp-EGFP* mouse brain descending from superficial motor cortex to deep hypothalamus across 6 mm of tissue. Volume size using of 1.5 × 1 × 6 mm (XYZ) acquired using a 5×/0.16 NA objective. Rendered using Imaris.

**Video 3. Myelin patterns in somatosensory cortex.** Fluorescent somatosensory cortex was isolated computationally from intact whole brain imaged with a 5×/0.16 NA objective. Volume size 4 × 4.4 × 4.6 mm (XYZ). Rendered using Imaris.

**Video 4. High resolution volume of layer 1 oligodendrocytes**. Volume from *Mobp-EGFP* brain acquired using high resolution 20×/1.0 NA objective. Volume size 1 × 0.5 × 1 mm (XYZ). Rendered using Imaris.

**Video 5. Whole spinal cord imaging of myelin patterns.** Whole-mounted fluorescent spinal cord from *Mobp-EGFP* mouse imaged with ZEISS lightsheet microscope using a 5×/0.16 NA objective. Volume size 4 × 27 × 3 mm (XYZ). Rendered using Imaris.

**Video 6. Mask R-CNN instance segmentation of individual oligodendrocytes in gray and white matter.** Cropped volumes (0.5 × 0.5 × 1.3 mm, XYZ) of gray and white matter from whole *Mobp-EGFP* brain (top). Mask R-CNN detected and segmented oligodendrocytes overlaid using multi-color masks (bottom).

## SUPPLEMENTAL INFORMATION

Document S1. Tissue processing and clearing guides, related to STAR Methods.

