## Supplemental Information (procedures) for "Brain-wide mapping of oligodendrocyte organization and oligodendrogenesis across the murine lifespan"

### Tissue Processing Guide

#### Tissue Preservation

*All steps done with shaking. Use 25mL tube, 1 per brain.*

| Solution | Amount | Time | Temp | Complete? |
|---|---|---|---|---|
| SHIELD OFF | 20mL* | 3 days | 4C | |
| SHIELD ON | 20mL** | ~1 day | 37C | |

**make fresh (day of)*
***prewarm solution in 37C water bath*

#### Clearing

*All steps done with shaking. Use 25mL tube, 1 per brain.*

| Solution | Amount | Time | Temp | Complete? |
|---|---|---|---|---|
| CUBIC L | 25mL | 3 days | 37C | |
| CUBIC L | 25mL | 3 days | 37C | |
| *CUBIC L** | 25mL | 3 days | 37C | |
| *CUBIC L** | 25mL | 3 days | 37C | |
| *CUBIC L** | 25mL | 3 days | 37C | |
| 20mM PB | 25mL | 4 hours | RT | |
| 20mM PB | 25mL | 2 hours | RT | |
| 20mM PB | 25mL | O/N | RT | |
| uRIMS | ~28mL*** | 2 days | 37C | |
| uRIMS*** | ~28mL | 1hr | 37C | |

**repeat as needed until clear (see table in long protocol)*
***fill 25mL tube to the top*
****if imaging in a different "stock" of RIMS, equilibrate in the new stock ~1 hour prior to imaging*

#### Mounting

- Use a UV-catalyzed glue (we like Ergo 8500) to adhere sample to holder
  - Add glue, then sample (not dripping), then glue.
  - Expose to UV light until hard (UV glue, ~15 sec).
    *Avoid drying out the brain as much as possible.*
- Incubate mounted sample in uRIMS for 15-60 mins at 37C prior to imaging.

### Materials and Solution Guide

#### SHIELD Preservation

SHIELD solutions are prepared in accordance with LifeCanvas's "Full Passive Pipeline Protocol" found on their website.

Reagents/Materials Needed:

- LifeCanvas™ Passive Tissue Clearing SHIELD Kit (SH-250 or SH-500)
- $dH_2O$
- Ice tray
- 25mL tube
- Curved forceps (for moving sample)

Steps:

1. Prepare SHIELD OFF fresh before each use by combining the following reagents **in order**. Per 1 mouse brain, combine:
   [5mL] $dH_2O$
   [5mL] SHIELD Buffer
   [10mL] SHIELD Epoxy
   Keep solutions on ice or move immediately to 4C. After each addition, vortex or mix well to prevent precipitation formation.

2. Before use, prewarm SHIELD ON solution (20ml/mouse brain) using a 37C water bath.

#### CUBIC L Delipidation

Delipidation is best achieved using the CUBIC L solution described in Susaki et al 2015. *Note:* CUBIC L will remove written markings on the sample tube – take care when handling the solution and labeling the tubes.

Reagents/Materials Needed:

- *n*-butyldiethanolamine (Tokyo Chemical Industry B0725)
- Triton x-100 (Sigma Aldrich T9284)
- $dH_2O$
- 1L bottle
- Stir bar
- Stir plate

Steps:

1. In a 1L bottle with a stir bar, combine:
   [100mL] *n*-butyldiethanolamine
   [100mL] Triton x-100
   [800mL] $dH_2O$
2. Mix on a stir plate for 30 mins or until fully combined. Do not shake the bottle to avoid the formation of large bubbles.

**Urea Refractive Index Matching Solution (uRIMS)**

Reagents/Materials:

- Dibasic sodium phosphate (Sigma Aldrich S9763)
- Monobasic sodium phosphate (Millipore Sigma 567545)
- dd$H_2O$
- Nycodenz (Serumwerk S18003) or Histodenz (Sigma Aldrich D2158)
- Diatrizoic acid (Sigma Aldrich D9268)
- Meglumine (Sigma Aldrich M9179)
- Sodium azide (Sigma Aldrich S2002)
- Urea (Sigma Aldrich U5378) or (Wako 210-01185)
- Hydrochloric acid (Fisher 210-01185)
- Triethylamine (Sigma Aldrich T0886)
- 500mL bottle
- 1L bottle
- Stir bar
- Large orbital shaker
- pH meter or pH strips
- refractometer

Steps:

1. <u>Make 20mM PB buffer.</u> In a 1L bottle with a stir bar, combine:
   [2.26g] Dibasic sodium. phosphate
   [0.46g] Monobasic sodium phosphate
   [1000mL] dd$H_2O$
   Mix until fully dissolved.
2. <u>Make uRIMS (~200mL).</u> In a 500mL bottle, combine:
   [160g] Nycodenz
   [2g] Meglumine
   [1.2g] Diatrizoic acid
   [0.016g] Sodium azide
   [88mL] PB buffer (20mM)
   Place the bottle on an orbital shaker at 37C and mix at 220rpm overnight or until fully dissolved.
   *Note*: the mixture will initially be too thick to use a stir bar or mix sufficiently by hand.
3. Once dissolved, add [110g] Urea to the bottle. Swirl to combine, then return the bottle to the shaker at 37C and mix to combine (6 hours-overnight).
   *Note*: the addition of urea produces an endothermic reaction.
4. The resulting solution with have a pH of 8-9. Use [~800µL] hydrochloric acid (HCl) to lower the pH to 7-7.5. If the pH becomes too acidic, use triethylamine (TEA) to bring it closer to neutral.
5. The final solution should have a refractive index ~1.48. Use a refractometer to measure the refractive index prior to imaging.
